# Characterization of Chemoresistant Cell Populations Improves Risk Stratification and Therapy Prediction in Pediatric AML

**DOI:** 10.1101/2025.09.25.678688

**Authors:** Mohammad Javad NajafPanah, Alexandra M Stevens, Michael J Krueger, Max Rochette, Sohani Sandhu, Lana Kim, Hua-Sheng Chiu, Jessica Epps, Sonal Somvanshi, Barry Zorman, Maria Rodriguez Martinez, Marianna Rapsomaniki, Susanne Unger, Burkhard Becher, Joanna S. Yi, Tsz-Kwong Man, Michele S Redell, Pavel Sumazin

## Abstract

Most pediatric acute myeloid leukemia (pAML) patients achieve complete remission after chemotherapy, yet relapse is common, with nearly 40% ultimately dying of the disease. Prognosis is currently assessed using cytogenetic biomarkers and measurable residual disease after the first chemotherapy cycle, with the highest risk patients referred for stem cell transplantation (SCT) at first remission. Because aggressive therapies such as SCT are highly toxic, yet cures after relapse are rare, accurate early risk prediction is essential for improving outcomes. To address this need, we analyzed paired diagnosis–relapse samples from 33 pAML patients at single-cell resolution and identified chemoresistant cell populations whose abundance at diagnosis significantly improved risk prediction. Incorporating the detection of these cell populations into our risk model revealed a previously unrecognized patient subgroup with a 5-year event-free survival rate below 40%. Although this subgroup represents only 20% of pAML cases, it accounted for half of the deaths among patients who do not receive SCT at first remission. Moreover, molecular characterization of these chemoresistant cell populations uncovered potential therapeutic targets and candidate interventions relevant to most high-risk patients, paving the way for more effective targeted treatments for high-risk pAML patients.

## INTRODUCTION

Over 40% of pediatric acute myeloid leukemia (pAML) patients relapse, including nearly 30% who are initially classified as low risk, and these relapsed patients have poor outcomes.^1–3^ Aggressive treatment options—such as high-dose chemotherapy, targeted therapies, and stem cell transplantation (SCT)—are available for high-risk patients and for those who relapse. Current methods for predicting relapse risk rely on identifying recurrent prognostic genetic variants at diagnosis and detecting measurable residual disease (MRD) after the first round of chemotherapy.^1,4^ SCT is frequently recommended for patients with high-risk variants or positive MRD.^5,6^ However, up to 30% of patients who test negative for MRD and do not carry high-risk variants still relapse.^3,7^ Early identification of these patients—before relapse, when SCT is most effective—is therefore critical for improving outcomes.^8^

A critical barrier to relapse prediction is the high intratumoral heterogeneity of pAML, in which individual patients often harbor leukemia cells with diverse DNA alterations, gene expression signatures, protein markers, and treatment responses. While some cell populations are effectively eradicated by standard chemotherapy, others can persist at essentially undetectable levels after treatment. Consequently, comprehensive molecular and clinical characterization of each cellular subpopulation within each leukemia is necessary to predict therapeutic response and evaluate relapse risk.^3,9^

Past efforts to characterize cellular subpopulations in rare pediatric and adult AML leveraged high-resolution RNA and protein profiling technologies.^3,10^ For example, single-cell RNA sequencing (scRNA-seq) profiles of adult AML patients revealed malignant cells with monocyte-like signatures that expressed T cell suppressing genes,^11^ and scRNA-seq analysis of chronic myeloid leukemia samples identified a response-predictive signature.^12^ Another single-cell profiling study of five paired AML samples collected at diagnosis and relapse identified common pathways associated with disease progression, including enhanced fatty acid oxidation and amino acid metabolism.^13^ While these studies provided insight into the biology of pAML, they neither improved current pAML risk prediction nor proposed new therapeutic strategies for high-risk pAML.^3^

Our prior work suggested that knowledge about the abundance of transcriptionally distinct pAML cell populations (*pAML aggregates*) and their response to therapies could help identify high-risk subpopulations.^10^ Towards this goal, we analyzed scRNA-seq expression profiles of paired diagnosis–relapse pAML samples obtained from 33 pAML patients together with healthy donor bone marrow aspirates. Here we report our analysis, which identified pre-treatment pAML aggregates that were intrinsically chemoresistant or acquired chemoresistance following therapy. We showed that their detection can both inform risk prediction in pAML clinical trials and predict the response of patient-derived xenograft models (PDXs) to chemotherapy. We also showed that the combination of chemoresistant pAML aggregate detection with cytogenetic risk models and MRD significantly improves risk prediction, revealing new high-risk and low-risk patient populations that may benefit from more and less aggressive therapies, respectively. Moreover, the detection of pAML aggregates that acquire chemoresistance during therapy revealed potential drivers of chemoresistance and vulnerabilities in these cell populations. We showed that targeting these inferred vulnerabilities using biological perturbations and small-molecule inhibitors in cell lines and PDXs specifically altered and depleted the targeted chemoresistant pAML aggregates.

## RESULTS

### Patient-focused analysis of longitudinal single-cell profiles

Because of the heterogeneous response of pAML subpopulations to therapies, we used longitudinal scRNA-seq to identify cell populations that were present before and after chemotherapy, as well as pAML subpopulations with evidence for chemoresistance acquisition. We analyzed paired diagnostic and relapse bone marrow samples from 33 pAML patients and six samples from non-cancer donors (Table S1). This included 22 Children’s Oncology Group (COG) AAML1031 patients from the Lambo cohort,^3^ four non-cancer donors profiled by Bailur et al.,^14^ and new paired profiles of 13 pAML patients plus two non-cancer donors—the Baylor College of Medicine (BCM) cohort. The BCM cohort included six Texas Children’s Hospital pAML patients, five previously uncharacterized COG AAML1031 patients, and two re-profiled COG cases that were included in the Lambo cohort. These re-profiled patients allowed for cross-cohort comparisons and reproducibility assessments.

As expected, longitudinal samples from the same patient were more transcriptionally similar to each other than to samples from different patients (Figures 1A,B). Surprisingly, even the BCM re-profiled diagnostic samples were more similar to their matched relapse samples than to the Lambo profiles of the same specimens (PAUVAT samples S3 and S53 and PAVBFN samples S11 and S55 in Figure 1). In contrast, non-malignant hematopoietic cell populations, including T cells and B cells, showed relatively similar transcriptomes across patients (Figure 1C; Table S2). Using a variety of representations of samples and their cellular composition, we observed both differences and similarities across cohorts, patients, time points, and cell types (Figure 1). These findings revealed that patient identity was the dominant driver of observed transcriptomic states, indicating that a patient-by-patient analytical strategy is appropriate (Tables S3–4).

**Figure 1.**
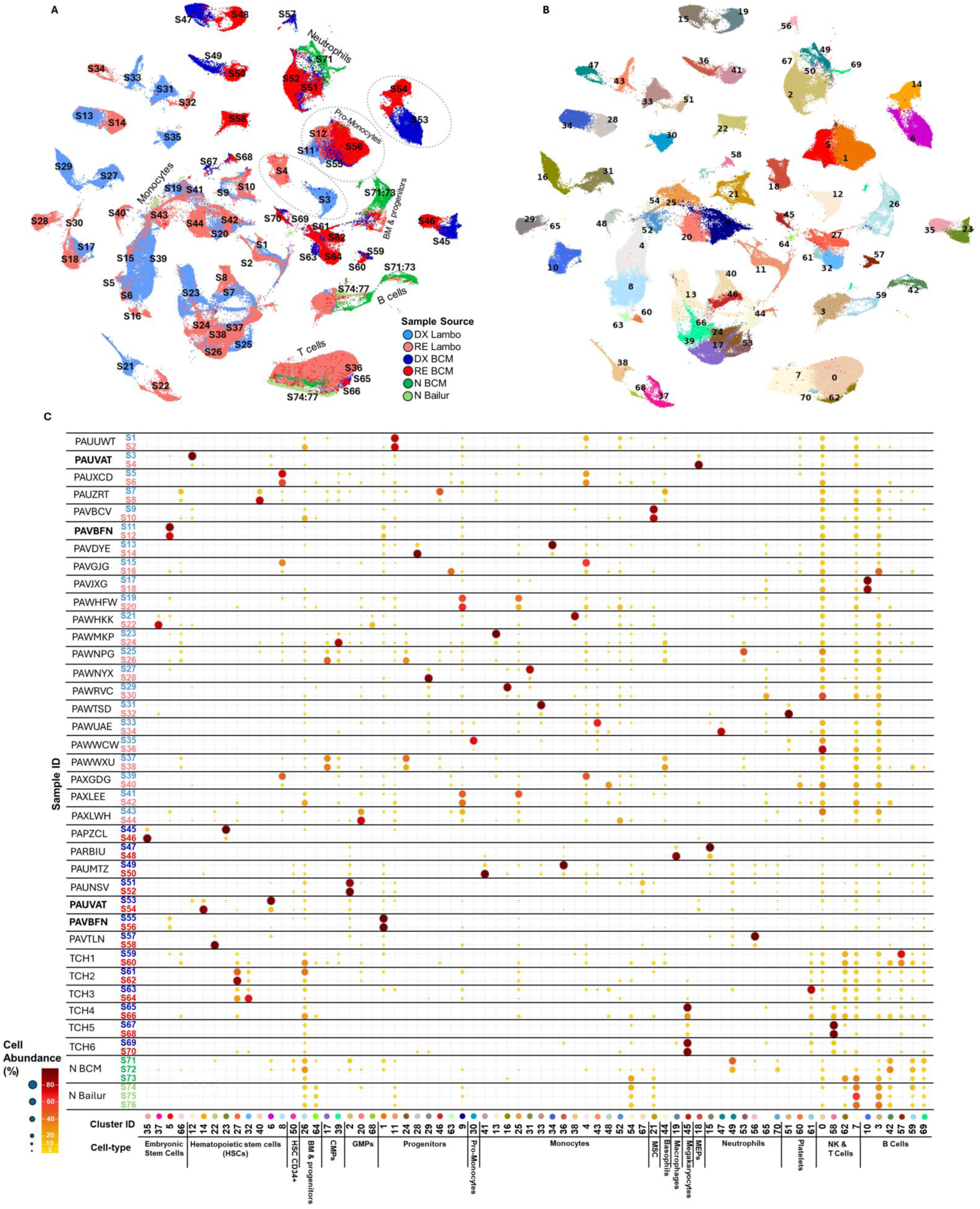
Merged single-cell RNA sequencing (scRNA-seq) profiles in pediatric acute myeloid leukemia (pAML) patients. **(A)** Uniform manifold approximation and projection (UMAP) visualizations of merged scRNA-seq expression profiles from paired diagnosis and relapse samples profiled by Lambo et al. (DX Lambo and RE Lambo) and this study (DX BCM and RE BCM), together with bone marrow profiles from healthy donors from this study (N BCM) and from Bailur et al. (N Bailur). (**B**) The clustering of scRNA-seq data by transcriptome similarity. **(C)** A comprehensive schematic summary of the identified cell clusters matching the UMAP in Figure 1B and indicating their inferred cell types and differentiation stages (columns), and sample sources with sample ID written using the color palette in Figure 1A (rows). Clusters were annotated using SingleR-inferred cell types. To aid visualization, relative cell abundance was recorded using both color and marker size. Samples from patients PAUVAT and PAVBFN were profiled in both the BCM and Lambo cohorts; these were circled (Figure 1A) and bolded (Figure 1C). GMP: Granulocyte-monocyte progenitor cells; CMP: Common myeloid progenitor cells; MEP: Megakaryocyte erythroid progenitor cells.

### Chemoresistant cells were inferred in most pAML cases

Using our patient-by-patient approach, we sought to address a critical but unresolved question in the field surrounding the etiology of chemoresistance in relapsed pAML patients: whether it is intrinsic or acquired. To distinguish intrinsic from acquired resistance, we developed a progression model describing the temporal dynamics of non-malignant, chemosensitive, and chemoresistant cell populations before and after therapy (Figure S1). This model delineates the expected presentation of non-malignant cells, chemosensitive pAML aggregates, and chemoresistant pAML aggregates in the profiles of patient samples before and after treatment.

Across all paired diagnosis–relapse samples, we identified 83 and 178 pAML *aggregates* in the BCM and Lambo cohorts, respectively. To align with our progression model, we classified pAML aggregates detected at diagnosis as *expanded*, *stable*, or *diminished pAMLs*, based on their pre-to post-therapy abundance ratios. Additionally, pAML aggregates detected only at relapse were termed *relapse pAMLs*. Expanded pAMLs were interpreted as intrinsically chemoresistant, while diminished pAMLs with evidence for transforming into relapse pAMLs were inferred to have acquired resistance.

Applying this model, we identified intrinsically chemoresistant pAML aggregates in 14 of 33 patients (42%; Group A, Figures 2A,B). The remaining patients were classified as Group B. This classification was corroborated by single-cell protein expression profiling by time-of-flight cytometry (CyTOF) in five paired cases (three Group A and two Group B; Figure S2) and remained consistent across the two re-profiled samples despite their technical variation. Notably, time to relapse did not significantly differ between the groups (Figure 2C). In addition, we identified diminished pAMLs that may have transformed into relapse pAMLs using trajectory analysis. This analysis supports transcriptomic transformations between adjacent cell populations across longitudinal samples from each patient (Figures 2D; S3). We identified diminished pAMLs with significant evidence for acquired chemoresistance in 19 of the 33 profiled pAML patients (58% of the pAML cases).

**Figure 2.**
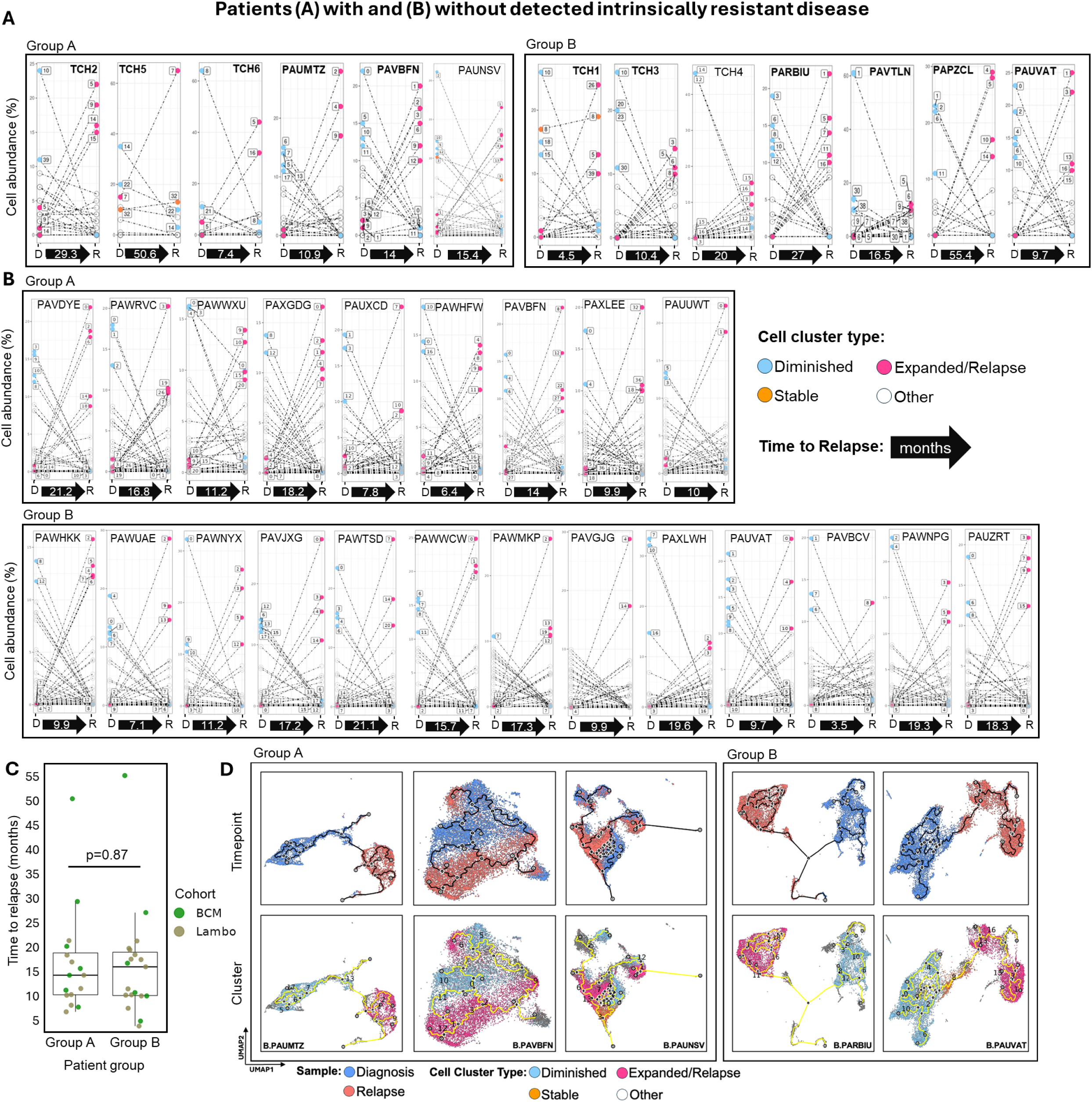
Longitudinal single-cell profiles identify patients with expanded and transforming pAMLs. Patients from the (**A**) BCM cohort and (**B**) Lambo cohort were classified based on the presence (Group A) or absence (Group B) of expanded pAMLs (intrinsically chemoresistant cell populations). Line plots map the estimated relative abundance of cell clusters identified at diagnosis and relapse; these were categorized into 4 cell types: Diminished, Stable, Expanded, and Other. **(C)** A comparison of the time to relapse for Group A versus Group B patients. **(D)** Trajectory plots based on scRNA-seq profiles of representative pAML pairs from Group A and B patients. UMAP color gradients represent timepoint (top) and transcriptome-based clustering (bottom). Predicted trajectories are illustrated with solid lines.

In total, chemoresistant pAML aggregates were identified in 32 of the 33 pAML cases. Of these, one case (PAUMTZ) was inferred to have both acquired and intrinsically chemoresistant pAML aggregates. Importantly, our analysis revealed that while acquired chemoresistance is common in pAML, nearly half of the pAML diagnosis samples also contained intrinsically chemoresistant cells. However, the abundance of these intrinsically chemoresistant cells at diagnosis was relatively low, accounting for 5-6% of the pAML cells in each sample, on average.

### Gene signatures are insufficient for predicting patient outcomes

Having inferred chemoresistant pAML aggregates in 32 of the 33 pAML cases, we asked whether their transcriptomic signatures could help predict patient outcomes. Towards this goal, we evaluated previously proposed outcomes-predictive pathways and gene sets^15–19^ and identified genes that were recurrently dysregulated in chemoresistant pAML aggregates (*Expanded Genes*, Tables S5,6). Multiple previously proposed gene signatures were upregulated in intrinsically chemoresistant cells, including genes associated with FMS-related receptor tyrosine kinase 3 (*FLT3*) and cyclin-dependent kinase 6 (*CDK6*) activation,^20^ the stem cell signatures LSC47 and HSC,^3,17^ and the MYC and oxidative phosphorylation pathways. As proposed by Lambo et al., this analysis confirmed an association between pAML chemoresistance and early hematopoietic embryonic differentiation stages (Figure S4).

To evaluate the potential of these gene signatures as prognostic biomarkers, we tested their outcomes-predictive ability using clinical and molecular profiles of AAML1031 patients produced by Therapeutically Applicable Research to Generate Effective Treatments (TARGET). We elected to focus on AAML1031 patients with no detected FLT3-ITD mutations to allow for treatment-independent patient outcomes comparisons. We found that all selected gene signatures could distinguish between diagnostic and relapse AAML1031 samples (Figure S4), but gene set enrichment analysis (GSEA) failed to predict outcomes for most pAML patients (Figure S5). Consistent with prior work,^3^ stem-cell signatures were predictive only in patients with KMT2A rearrangements. We reasoned that because intrinsically chemoresistant pAML aggregates represented only 5–6% of the pAML cells in diagnostic samples, their signal may be too weak to be reliably detected by GSEA.

### Chemoresistant pAML aggregate abundance improves risk prediction

Our prior work suggested that the impact of low-abundance chemoresistant pAML aggregates could be detected in bulk RNA profiles using deconvolution.^9^ We therefore tested whether the estimated abundance of chemoresistant pAML aggregates is predictive of AAML1031 patient outcomes. In total, we identified five BCM-and eighteen Lambo-cohort chemoresistant pAML aggregates whose inferred abundance each significantly predicted both overall survival (OS) and event-free survival (EFS) of AAML1031 patients (adjusted p<0.05; *predictive aggregates*). Of these, four and seven aggregates were inferred to acquire chemoresistance in the BCM and Lambo cohorts, respectively. To minimize the analytical uncertainties introduced by integrating these two cohorts, we elected to infer the abundance of predictive aggregates in parallel. As expected, the inference of their total abundance—calculated as the sum of the abundances of predictive aggregates in each dataset—was significantly more predictive of OS and EFS than that of any individual predictive aggregate (p<1E-4 by F-test), and the combination of the two quotients significantly improved on individual predictive abilities.

Importantly, predictive aggregate abundance improved risk prediction in AAML1031 patients beyond cytogenetics and further refined MRD-based stratification. To evaluate the benefit of predictive aggregate abundance, we tested whether it could improve on the risk-prediction framework proposed in the ongoing COG AAML1831 trial (Figures 3A; S6A). Validation in two independent cohorts, AML08^9^ and AAML0531^21^ (Figures 3B; S6B), confirmed that predictive abundance enhanced outcome classification without additional model fitting (p = 1E-4 for both EFS and OS; Table S7). While improvements were evident across nearly all risk categories, we focused on those with the greatest potential to affect treatment decisions in COG AAML1831.

**Figure 3.**
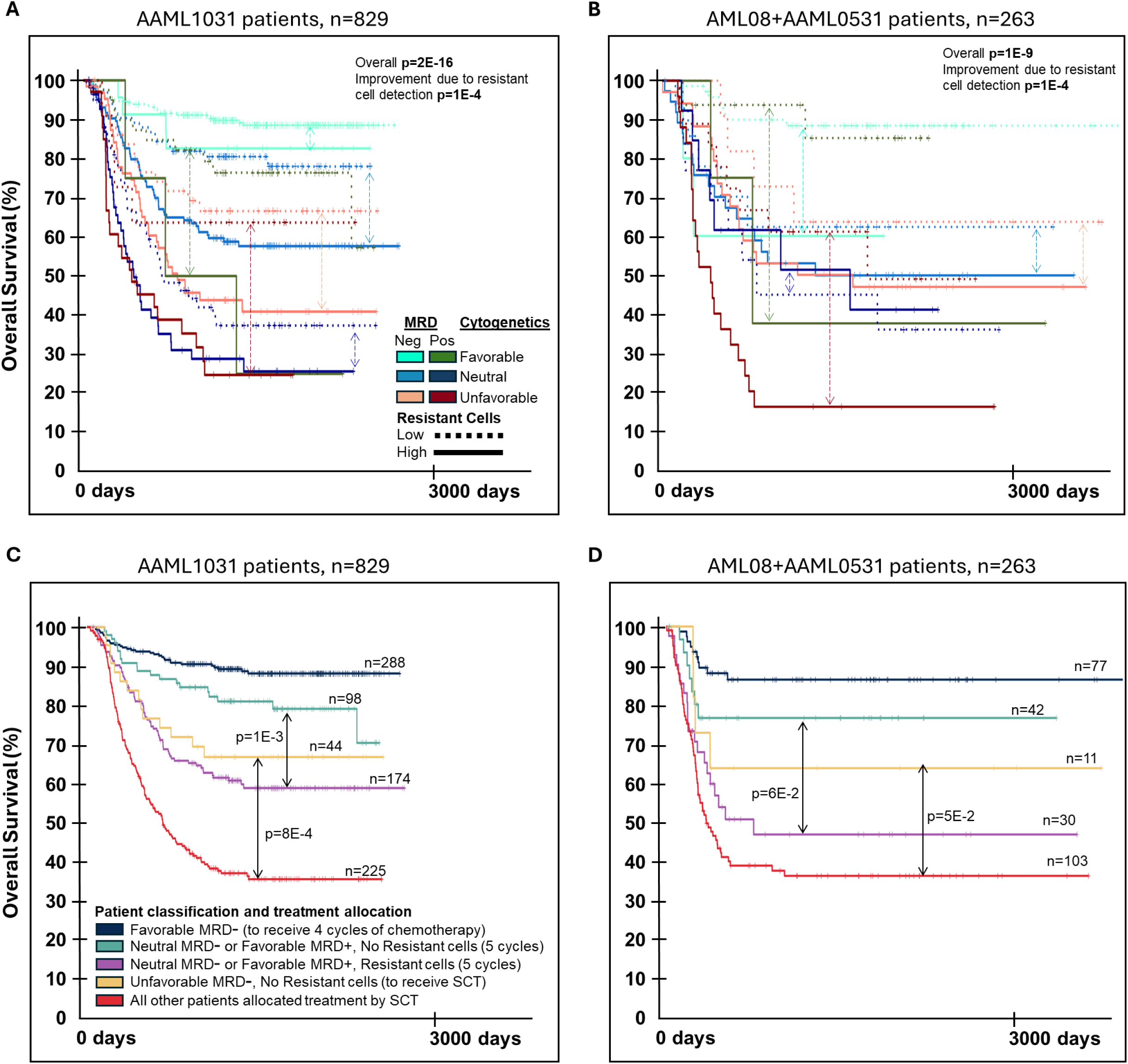
Overall survival analysis of predictive pAML aggregates. (**A–B**) Estimated abundance of chemoresistant pAML aggregates (resistant cells) significantly predicted patient outcomes in (A) AAML1031 and (B) AML08 + AAML0531 cohorts, as determined by Kaplan– Meier survival analysis. Patients were stratified by MRD (darker shading indicates positive) after one cycle of chemotherapy and stratified by cytogenetics according to the COG AAML1831 protocol to Favorable, Neutral, or Unfavorable (color-coded green, blue, and red, respectively). Across independent trials, resistant-cell abundance improved OS prediction (p = 1E-4). Solid and dotted lines represent high-and low-resistant-cell-abundance patients, respectively. Arrows indicate patients within the same AAML1831 risk categories. (**C–D**) Resistant cell abundance reclassified patients in (C) AAML1031 and (D) AML08 + AAML0531 who would be allocated identical therapies under the current AAML1831 protocol. This analysis revealed higher-risk patients (purple) not prescribed SCT under the COG AAML1831 protocol, with OS < 0.6 and EFS < 0.4 (Figure S6), and lower-risk patients (orange) who would be prescribed SCT but would achieve OS > 0.6 and EFS > 0.4 without SCT across AAML1031 and AML08 + AAML0531.

Under the AAML1831 protocol, patients with Neutral cytogenetics and negative MRD and those with Favorable cytogenetics and positive MRD (LR2 patients) receive five rounds of chemotherapy, whereas patients with Unfavorable cytogenetics and negative MRD (HR patients) are allocated SCT. We retrospectively applied the AAML1831 risk group definitions to the AAML1031, AAML0531, and AML08 cohorts, recognizing that as the definition of High Risk evolved, so too did the characteristics of patients who did and did not undergo SCT at first remission. Our analysis showed that LR2 patients with a high predictive aggregate abundance (LR2-high) had significantly worse OS and EFS than LR2-low patients (Figures 3C,D; S6C,D). The 5-year EFS for the LR2-high patient category in AAML1031 and AML08+AAML0531 were 38% and 27%, respectively. Strikingly, LR2-high patients represented ∼20% of each cohort but accounted for ∼50% of deaths among patients not prescribed SCT. Conversely, although fewer patients were classified as HR with low predictive aggregate abundance (HR-low), their EFS (>40% across trials) and OS (>64% across trials) were dramatically better than those of other SCT-allocated patients (EFS<30% and OS<40%; p=5E-4 for AAML1031). Notably, while most patients who would be prescribed SCT in AAML1831 but not in prior trials died of disease, two-thirds of HR-low patients survived without SCT across the tested cohorts (Table S7).

### Predictive aggregates stratify outcomes across cytogenetic classes

Given that the detection of predictive aggregates significantly improved risk prediction after consideration of both cytogenetic risk and MRD, we next explored whether their predictive ability was linked to their cytogenetic subtype. Namely, we evaluated the relationship between predictive aggregates, the cytogenetic classification of the patients from which these predictive aggregates were derived, and the cytogenetic classification of the patients whose outcomes they predicted (Figure S7). Because KMT2A rearrangements were frequent in the BCM and Lambo cohorts, and due to the complexity of the AAML1831 protocol, we elected to investigate the traditional cytogenetic categories employed by Lambo et al. (2023); a mapping between these cytogenetic categories and the three risk categories used in AAML1831 across trials is given in Figure S7C. The cytogenetic categories that were most strongly associated with poor pAML patient outcomes were *KMT2A* Rearrangement and Other. Our analysis revealed that most predictive aggregates were identified in samples from these high-risk pAML cytogenetic categories. Only three (13%) were derived from lower-risk cases, including PAVGJG (inv(16)), and PAWMKP and PAWNPG (t(8;21))—all Lambo cohort cases. We note, however, that patients with the highest risk cytogenetics also accounted for most of the BCM and Lambo cohorts, 64% and 73%, respectively.

Notably, predictive aggregates that were identified in samples from high-risk pAML cytogenetic categories effectively classified the outcomes of patients with low-risk cytogenetics (Figures 3A,C,D). For example, while no predictive aggregates were sourced from patients in the Normal cytogenetics category, predictive aggregates significantly improved outcomes prediction for both MRD-positive and MRD-negative patients with Normal cytogenetics (Figure S7). However, predictive aggregates were significantly more abundant in patients whose cytogenetic category matched the source patients, e.g., predictive aggregates characterized in samples from KMT2A patients were enriched in other KMT2A patients (Figure S7). This suggests that while predictive aggregates were best at predicting outcomes for patients with similar cytogenetic backgrounds, aggregates derived from samples with high-risk cytogenetic biomarkers can also be predictive of the outcomes of patients identified as low risk by cytogenetics.

### Chemoresistant pAML cell abundance predicts clonal PDX response to chemotherapy

Having demonstrated that predictive aggregates selected based on their fit with our model for chemoresistance were predictive of clinical outcomes in retrospective analysis of multiple clinical trials, we directly tested whether they were also predictive of chemoresistance using pAML PDXs.

Towards this goal, we evaluated tumor responses to cytarabine in eight models from the Pediatric Acute Leukemia Xenograft (PALeX) resource.^22^ We compared the proportion of human pAML cells in PDX bone marrow and spleen after treatment for 4 days with cytarabine to PDXs treated with saline (Figures 4A; S8, and Table S8). All PDXs were derived from samples of pAML patients who were treated with cytarabine-containing regimens. We identified two PDXs with significant sensitivity to cytarabine (AML006 and AML005). Of the remaining six PDXs, two (AML001 and AML010) showed some response to cytarabine, but this response was not statistically significant. The remaining four, including AML903 and AML905, showed no indication of sensitivity to cytarabine (Figure 4B).

**Figure 4.**
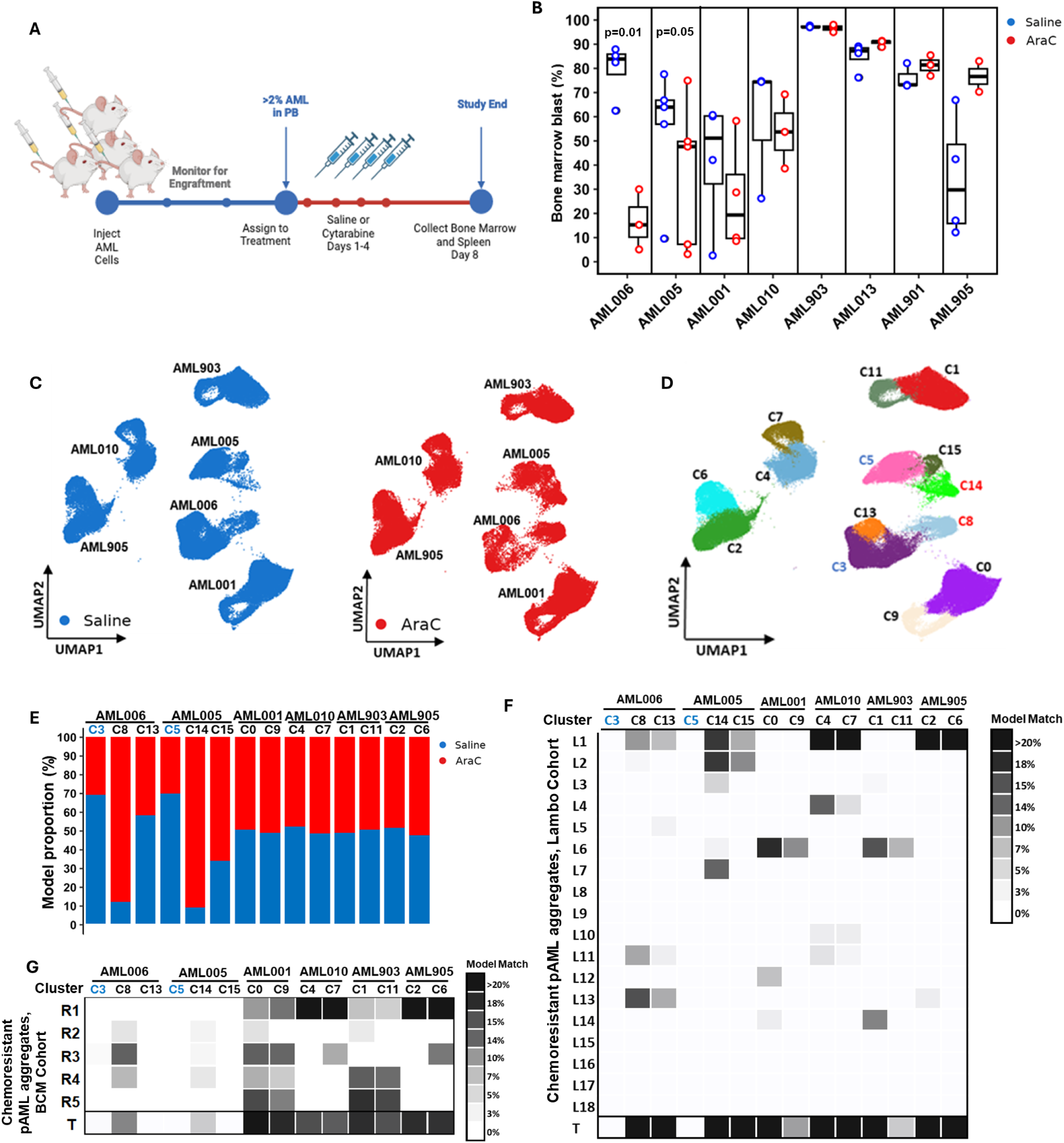
**Outcome-predictive pAML aggregates predict PDX chemotherapy response**. (**A**) PDX models with established tumor burden were treated with 4-day courses of cytarabine (AraC) or saline and then euthanized for disease evaluation. (**B**) Bone marrow blast proportions revealed significant AraC sensitivity in AML006 and AML005. (**C**) scRNA-seq profiles of saline-and AraC-treated AML006, AML005, and four less chemosensitive PDXs revealed (**D)** PDX-specific, but not treatment-specific, transcriptomes, with 2–3 distinct clusters per model. (**E**) Cluster-level analysis revealed AraC-sensitive clusters (C3, C5) and AraC-resistant clusters (C8, C14) in AML006 and AML005, respectively, while other PDXs showed no treatment-related cluster-abundance differences. (**F–G**) Deconvolution mapped PDX clusters to predictive pAML aggregates from the (F) Lambo and (G) BCM cohorts. AraC-resistant clusters C8 and C14 matched inferred chemoresistant pAML aggregates in patients, whereas the AraC-sensitive clusters C3 and C5 did not.

To evaluate clonal response to chemotherapy, we profiled representative AML006, AML005, AML001, AML010, AML903, and AML905 PDXs (two from each response type) by scRNA-seq. Consistent with our observations in human tumors, scRNA-seq profiles of cytarabine-treated and saline-treated PDXs revealed PDX-specific molecular signatures (Figure 4C), with two to three distinct transcriptionally-defined cell clusters identified per model (Figure 4D; Table S9). Our analysis revealed expanding and diminishing pAML cell populations in the chemosensitive AML006 and AML005 but not in other PDXs, in which the abundance ratios after treatment by cytarabine and saline were unchanged (Figure 4E). Namely, cells from clusters C3 and C5 accounted for 77% and 87% of the saline-treated pAML cells in AML006 and AML005, respectively, but these clusters showed significantly lower abundance in cytarabine-treated mice (p=1E-3 by student’s *t*-test, Table S9). In contrast, cells from clusters C8 (AML006) and C14 (AML005) had a 3-fold increase and a significantly higher abundance in cytarabine-treated models (p=1E-4, Student’s *t*-test). Thus, we concluded that the abundance of C3 and C5 cells decreased after treatment with cytarabine due to sensitivity to chemotherapy, while C8 and C14 cells increased due to chemoresistance. Analogously, this analysis suggested that pAML cells that were characterized in other PDXs were chemoresistant.

The identification of chemosensitive and chemoresistant PDX pAML cell populations allowed us to evaluate the correspondence between predictive pAML aggregate abundance and cytarabine response.^23^ Towards this goal, we evaluated how well each PDX pAML cell cluster transcriptomically matched predictive aggregates identified in the Lambo (Figure 4F) and BCM (Figure 4G) cohorts. Our analysis revealed that patient-derived predictive aggregates—from both the BCM and Lambo cohorts—exhibited a high degree of transcriptional similarity to pAML cells in all clusters identified as chemoresistant (Table S10). However, this similarity was not observed in any cells from the chemosensitive clusters C3 (AML006) and C5 (AML005). This result directly supported our assertion that our predictive aggregates represent bona fide chemoresistant cell populations.

### Targeting pathways associated with chemoresistance acquisition

The identification of patient pAML aggregates that were inferred to acquire chemoresistance presented an opportunity to infer molecular mechanisms that may drive chemoresistance and to identify vulnerabilities in these cells. We identified dysregulated gene sets and regulatory modules—based on both differential RNA expression and chromatin accessibility across pAML aggregates from the same diagnostic sample—that were enriched in diminished pAMLs that were inferred to acquire chemoresistance (Figure S9). These included pAML signatures that were tested for risk prediction improvement for AAML1031 patients (Figure S4) and regulatory network modules downstream from transcription factors, as identified in our pAML-specific reverse-engineered regulatory network.^24–26^ This analysis identified recurrent dysregulation in FLT3-and CDK6-associated gene sets, as well as pathways downstream from or associated with RXRA and RARA, MYC, G2–M checkpoint, p53, TNFα, KRAS, and ALK.

To test the therapeutic relevance of targeting these gene sets and pathways, we curated publicly available data from assays that targeted inferred vulnerabilities in pAML cells and tumors. These data included assays using biological and small molecule inhibitors to target IRF8, CHAF1B, KLF4, DNMT1, BCL6, CDK4/66, and ATF3 (Figures 5A-F and S10A-F).^27–34^ We also used small molecule inhibitors to target FLT3, CDK4/6, and RARA in patient-derived pAML cells in vitro and pAML PDXs in vivo (Figures 5G-K and S10G-K). For both the curated data and our own experiments, we compared the observed changes—comparing targeting agents and controls—in the abundance of targeted pAML aggregates versus non-targeted control pAML aggregates. Across all perturbations, targeted pAML aggregates showed consistent depletion upon targeted inhibition, while pAML aggregates that were detected in the same samples but were not predicted to respond to the specific agent were unaffected by the treatment (Figures 5A–L, S10). Moreover, 60% of AAML1031, AML08, and AAML0531 patients who would be allocated SCT in COG AAML1831 had a high abundance (>10%) of at least one targeted predictive aggregate with significant response to one of the five tested inhibitors (Figure S11; Table S13). This analysis supports the therapeutic potential of these inhibitors for high-risk pAML patients, including FLT3 inhibitors like quizartinib, which also inhibits the wild-type FLT3, even for FLT3-ITD-negative patients.^35^

**Figure 5.**
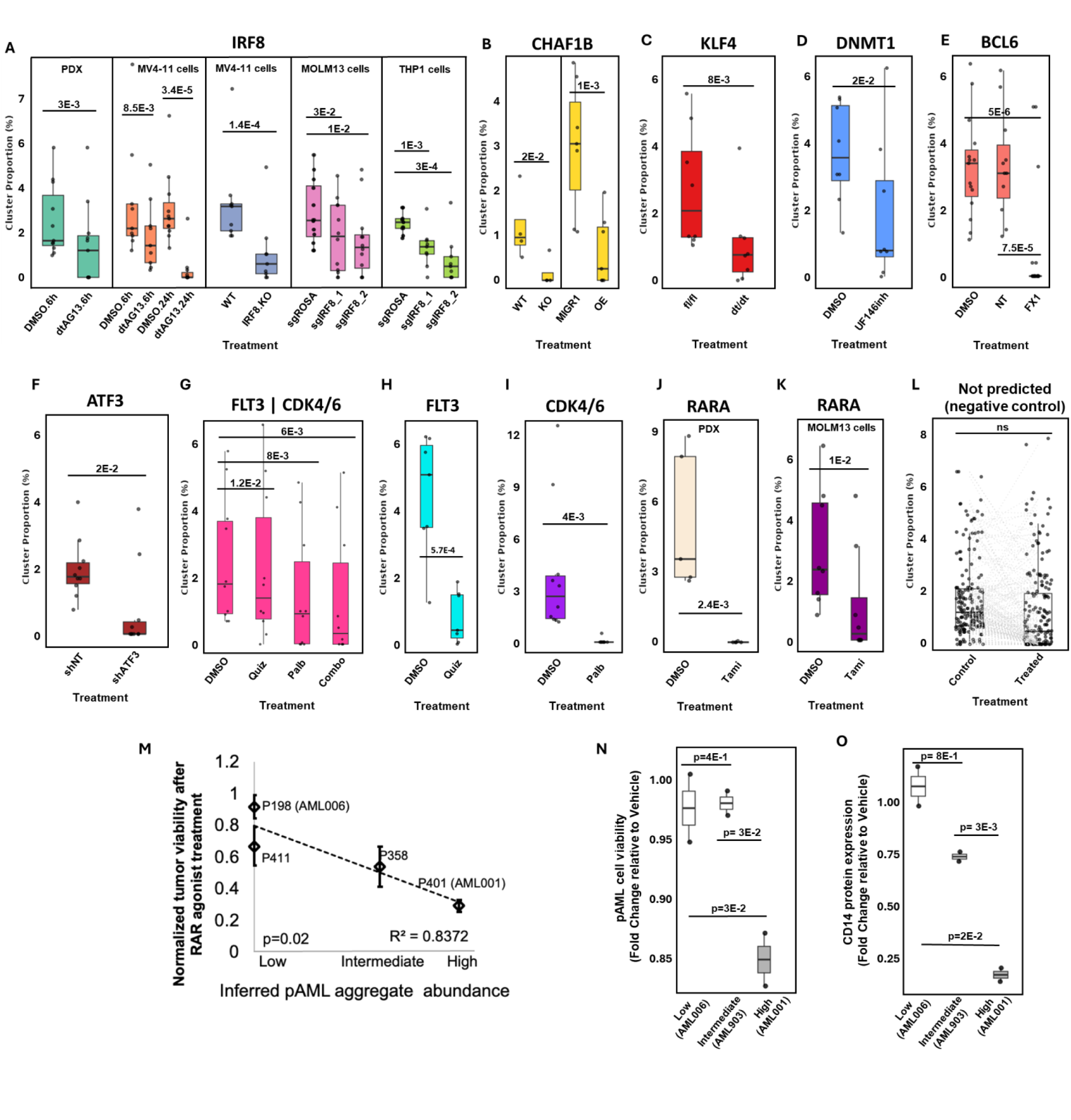
Predicted vulnerabilities in chemoresistant pAML aggregates. (**A–K**) Predicted vulnerabilities were validated using public perturbation datasets (A–F) and de novo assays (G– K), showing significant declines in the abundance of targeted pAML aggregates after treatment with biologicals or small-molecule inhibitors. The pathway or gene being tested is listed above each panel, and the control condition is shown on the left. (**L**) Aggregates not predicted to be vulnerable showed no significant changes; see Figure S10 for details. (**M**) Predicted vulnerability to the retinoic acid receptor agonist tamibarotene correlated with tumor cell viability in four pAML patient samples; note that samples from patients p198 and p401 were used to generate PDXs AML006 and AML001. (**N–O**) Predicted vulnerability to the FLT3 inhibitor quizartinib and CDK4/6 inhibitor palbociclib was confirmed by (N) reduced cell viability and (O) CD14 protein expression relative to DMSO controls (mean ± s.e.m.). DMSO: dimethyl sulfoxide control; dtAG: degradation tag; KO: knockout; WT: wildtype; NT: non-targeting; fl/fl: floxed/floxed; dt/dt: degradation tag; OE: CHAF1B overexpression; UF146inh: UF146 inhibitor; Quiz: quizartinib; Palb: palbociclib; Tami: tamibarotene.

Next, we sought to confirm the impact of targeting these gene sets and pathways on tumor viability by testing whether the abundances of pAML aggregates that were inferred to be vulnerable to (1) retinoic acid receptor activation and (2) FLT3 and CDK4/6 inhibition are predictive of PDX responses to these treatments. Our RNA profiles of PDX models targeted with the second-generation retinoic acid receptor agonist tamibarotene by Perez et al.,^36^ revealed that the abundance of pAML aggregates with suppressed RXRA-and RARA-activity predicted response to tamibarotene. Specifically, the inferred total abundance of these pAML aggregates was significantly correlated with the observed change in tumor viability upon treatment with tamibarotene in vitro (R^2^=0.8, p=0.02, Figure 5M).^37^ Patient sample P401 and its corresponding PDX model AML001 had a high abundance of cells with suppressed retinoic acid activity and were highly sensitive to tamibarotene in vitro and in vivo. In contrast, patient sample P198 (PDX model AML006) and P411, lacking these pAML aggregates, did not show benefits (Figure 5M).

Analysis of scRNA-seq data of our PDX panel (Figures 4C-D) suggested that neither of the three clusters of the chemosensitive AML006 contained pAML aggregates that are enriched for the upregulation of FLT3-and CDK4/6-associated gene sets. In contrast, one of the clusters of the chemoresistant AML903 and both clusters of the chemoresistant AML001 contained such aggregates. This suggested that AML903 and AML001, but not AML006, may be vulnerable to the FLT3 inhibitor quizartinib and the CDK4/6 inhibitor palbociclib. Treatment of these PDX cells in vitro revealed that tumor viability and protein expression of CD14—a treatment-induced differentiation marker that is upregulated in the targeted predictive aggregates—significantly correlated with the abundance of pAML aggregates that were predicted to be vulnerable to FLT3 and CDK4/6 inhibition (Figures 5N,O). Treatment of AML903 in vivo with the combination of quizartinib and palbociclib increased median survival by 210%, from 29 to 61 days. Note that AML903 contained predictive pAML aggregates that responded to both quizartinib and palbociclib (Figures 5H,I and S10H,I). Taken together, these findings demonstrated that predictive aggregates not only stratify patient risk but also nominate actionable vulnerabilities that could be exploited to target chemoresistant pAML cell populations.

## DISCUSSION

Guided by a model of pAML progression (Figure S1), we analyzed longitudinal scRNA-seq profiles of paired pAML samples collected before and after chemotherapy to identify and characterize pAML aggregates that are intrinsically chemoresistant or that acquire chemoresistance following treatment. Our results indicated that chemoresistant cells were already present in diagnostic samples from approximately half of the patients who eventually relapsed, although they represented only 5-6% of the total pAML population at diagnosis (Figure 2). This finding underscores the need for technologies capable of detecting rare, low-abundance chemoresistant cells prior to relapse.

To this end, we applied the deconvolution method SQUID^10^ to RNA-seq profiles of patients enrolled in the AAML1031, AML08, and AAML0531 clinical trials. Our analysis of patient data from these clinical trials suggested that the presence of scRNA-seq-characterized chemoresistant pAML aggregates is an effective classifier for both MRD-negative patients with high-risk cytogenetics and MRD-positive patients with low-risk cytogenetics (Figures 3 and 4). Adoption of this improved risk-predictive approach has the potential to enable earlier identification of MRD-negative and additional MRD-positive patients who would benefit from aggressive treatment, including SCT and targeted therapies, before relapse, when outcomes from these treatments are most favorable. Such an adaptation of treatment protocols has the potential to improve survival for the 40% of pAML patients who currently experience poor outcomes.

In addition, we showed that molecular characterization of outcome-and chemoresistance-predictive aggregates can reveal their potential therapeutic vulnerabilities. Functional assays targeting IRF8, CHAF1B, KLF4, DNMT1, BCL6, FLT3, CDK4/6, RXRA, and ATF3 in cell lines and animal models showed that chemoresistant pAML aggregates predicted to be sensitive to a given agent were specifically and significantly altered, while other pAML aggregates were not affected (Figures 5 and S10). Collectively, these findings suggest that identifying and molecularly profiling chemoresistant pAML populations can both improve relapse prediction and inform the development and selection of personalized targeted therapies for high-risk patients.

The identification of predictive chemoresistant pAML aggregates confirmed the relevance of key pAML pathways. Yet, as previously observed,^3^ the enrichment of these gene sets in bulk profiles of diagnostic samples failed to predict outcomes for most patients. Namely, pathways associated with chemoresistance and hematopoiesis stemness, including MYC,^1^ FLT3, CDK6, the oxidative phosphorylation pathway,^38^ and the stem cell signatures LSC47 and HSC^3,17^, were significantly upregulated in predictive aggregates and relapse samples, but the corresponding gene sets were not significantly enriched in the diagnostic RNA-seq profiles of high-risk patients. Our analysis suggested that relapse may be driven by rare chemoresistant cell populations and that technologies to predict relapse need to detect signals produced by as few as 5% of the pAML cells. Methods akin to gene-set enrichment that rely on the dysregulation of predictive biomarkers in bulk profiles of pAML samples are ill-equipped to detect such rare populations. To overcome this obstacle, we leveraged computational technology developed in prior work, where we showed that RNA-seq deconvolution is sensitive enough to detect and estimate the abundance of rare cells.^10^

By applying this technology to predict outcomes for patients in the AAML1031, AML08, and AAML0531 trials, as well as to predict PDX responses to chemotherapy, we were able to identify chemoresistant pAML cells that drive relapse and patient outcomes. Our results rely on the integration of multiple independent lines of evidence. First, we developed a progression model to define the expected dynamics and distributions of pAML and non-malignant cells before and after treatment (Figure S1), enabling us to nominate candidate chemoresistant populations. We then tested these candidates for their ability to predict relapse and patient outcomes. To minimize overfitting, we applied only limited machine learning when training our predictive algorithm on molecular and clinical profiles of AAML1031 patients, and used no additional learning when assessing risk in the AML08 and AAML0531 cohorts. These analyses confirmed that our inferred chemoresistant pAML aggregates are predictive of patient outcomes. Finally, we validated their chemoresistant nature by showing that they are molecular matches to chemoresistant PDX populations, including rare resistant subpopulations within PDXs that are otherwise predominantly composed of chemosensitive pAML cells.

Current risk-prediction strategies combine cytogenetic analysis at diagnosis with MRD detection during treatment,^9,39^ and our analysis confirmed the strength of these predictors. In diagnostic samples from FLT3-ITD–negative AAML1031 patients, we observed that tumors with core binding factor fusions (CBFB::MYH11 and RUNX1::RUNX1T1) or normal karyotypes were associated with more favorable outcomes than those with KMT2A rearrangements or other cytogenetic abnormalities. While most of our predictive aggregates originated from these high-risk patients, we found that they also predicted outcomes among lower-risk patients, albeit at lower prevalence (Figure S7). This suggests that these aggregates encapsulate high-risk features that are independent of current cytogenetic biomarkers and risk stratification methods. Notably, integrating all three predictive approaches yielded significantly higher accuracy than any two in combination, underscoring their complementary strengths.

Our strategy for predicting patient outcomes, chemotherapy resistance, and response to targeted therapies centered on estimating the abundance of chemoresistant pAML aggregates. This builds naturally on existing approaches that assess relapse risk through MRD detection. However, instead of quantifying residual pAML cells after treatment, we estimated the abundance of chemoresistant pAML cells prior to treatment, as well as pAML populations with specific vulnerabilities prior to their therapeutic targeting. Importantly, we demonstrated the feasibility of this approach using both computational deconvolution of bulk profiles and more direct single-cell– resolution profiling. Notably, our analyses also suggested that the vulnerabilities we identified, along with potential drivers of pAML progression, are epigenetically regulated. In particular, we found that regulatory networks differentially activated in chemoresistant pAML cells are frequently controlled through chromatin accessibility (Figure S9).

Our current approach has several noteworthy limitations. First, our model does not account for the possibility that pAML cells may arise de novo from non-malignant cells following chemotherapy, or that convergent evolution may occur when chemoresistant cells adopt transcriptomic profiles resembling chemosensitive cells. Second, the model assumes that relapse pAML cells lacking pre-treatment chemosensitive counterparts are more likely to be chemoresistant. These assumptions may not always hold; however, the model serves only as a screening framework to select candidates, who are subsequently tested for their ability to predict patient outcomes before being established as predictive pAML aggregates. Finally, although we demonstrated that pAML aggregates with predicted vulnerabilities are altered by therapies targeting those vulnerabilities (Figures 5A–L), we did not directly establish that these therapies reduce the viability of the corresponding cells. Such treatments may instead induce transient transcriptomic state changes without compromising survival. To address this limitation, we evaluated PDX models treated with a RARA agonist and FLT3 and CDK4/6 inhibitors. In these experiments, PDXs enriched for predicted vulnerable populations showed significantly reduced viability (Figures 5M–O), supporting the conclusion that pAML aggregates with predicted vulnerabilities can indeed be selectively targeted. Note that prospective validation of our prognostic model can be achieved with bulk RNA-seq profiling of newly diagnosed patients who are followed for treatment response and outcomes.

## METHODS

### Patient samples

Bone marrow or blood samples were collected at diagnosis and relapse for pAML patients whose families consented to banking tissue for research in accordance with the guidelines of the Declaration of Helsinki. Six diagnosis–relapse pairs were obtained from patients diagnosed and treated at Texas Children’s Hospital (TCH). Seven diagnosis–relapse pairs were obtained from the COG Biopathology Center. All patients received initial chemotherapy while enrolled in or according to the protocol used in the Phase 3 COG clinical trial AAML1031^7^. Two normal bone marrow samples were obtained from healthy donors by rinsing the discarded collection filter. All COG samples were enriched for mononuclear cells by density centrifugation and cryopreserved; these samples were subjected to fluorescence-activated cell sorting (FACS) to enrich for AML cells (side scatter [SSC] low, CD45 dim) prior to downstream analyses. Viably frozen samples from the TCH cases were sent for single-cell sequencing without prior sorting. To control the effects of sorting, one normal bone marrow aspirate was split into two, and only half the cells were FACS sorted to enrich for immature myeloid cells in the same gate as used for the COG cases. All cases passed quality control (QC) for inclusion in our analysis. All paired diagnosis-relapse samples with clinical annotations from Lambo et al. were included in the study. When calculating EFS, events included induction failure, relapse, and death.

### Cytogenetic classification and treatment in the current COG trial

We followed the COG AAML1831 trial protocol for assigning risk based on cytogenetic biomarkers. COG AAML1831 is expected to accrue patients until 2030. The biomarkers used to classify patients into Favorable, Neutral, and Unfavorable cytogenetics categories are provided for readability in Table S7. We relied on published trial data to determine MRD, which was evaluated independently in each trial, with positive MRD identified at 0.1%. Treatment allocation in AAML1831 is as follows. Patients with Favorable cytogenetics and negative MRD are allocated 4 cycles of chemotherapy; patients with Favorable cytogenetics and positive MRD and patients with Neutral cytogenetics and negative MRD are allocated 5 cycles of chemotherapy; the remaining patients, including patients with Neutral cytogenetics and positive MRD and patients with Unfavorable cytogenetics, are allocated 3 cycles of chemotherapy followed by SCT.

### scRNA-seq profiling and analysis

#### Profiling

Patient samples were flow-sorted, gating for CD45-dim and SSC low cells; 10K–15K cells were then labeled using the 10x Genomics (Pleasanton, CA) Chromium Next GEM Single Cell 3’ Kit v3.1 and sequenced at 200 M reads per sample using an Illumina NovaSeq 6000. The resulting scRNA-seq data were analyzed by an anchor-based approach with the *Seurat* package v4.30. Specifically, cells were analyzed using three distinct methods. First, cell counts were log-normalized and clustered using the Louvain algorithm. Cells across all profiled and collected samples were then merged (Figure 1A) using standard *Seurat* v4.30 packages^40^. Second, cell profiles from paired samples and normal samples were normalized and integrated one patient at a time to classify pAML and non-tumor cells. Finally, patient-specific cell clusters were combined to merge similar cells within and across patients in an effort to reduce the complexity of the transcriptomic space.

#### Cell classification

To analyze patient-specific cell populations, we first log-normalized the count data from each patient pair containing diagnosis and relapse samples together with data from three normal bone marrow samples from two individuals (TCHnorm1 and TCHnorm2). Normal bone marrow samples were included to help distinguish malignant from non-malignant hematopoietic cells and to accommodate for the smaller number of cells associated with patient-by-patient analysis. Normalized data were then integrated by their sample types (tumor, normal) using the *IntegrateData* function, and cell clustering was performed on the integrated data using the Louvain algorithm at a resolution of 2.5. Clusters with fewer than 5% of the cells from each normal bone marrow sample were considered pAML candidates; inferred cell types were verified at a 100% rate as potential cancer cells. We identified expanded pAMLs as integrated cell clusters comprising cells from both the diagnosis and relapse samples. Expanded pAMLs had to include >10% of the cells from the relapse sample. Similarly, diminished pAMLs were identified as clusters including >10% of the cells from the diagnosis sample, whereas stable pAMLs were classified as those for which >10% of cells from both paired samples passed QC. Clusters that included >10% of the cells from the relapse sample only were identified as relapse pAMLs. The remaining clusters, including small clusters or clusters that were mostly composed of non-pAML cells, were identified as *other cells*. When combining clusters across patients, we merged clusters with similar (pseudobulk) expression profiles and argued, based on parameters set by SQUID^10^, that these clusters are derived from the same pAML aggregates.

#### Estimating cell population abundance based on bulk RNA-seq profiles

We used the SQUID framework to predict the abundance of scRNA-seq clusters and pAML aggregates in bulk RNA-seq profiled samples.^10^ The first step of SQUID’s framework is the combination of cell clusters with similar expression patterns. This is accomplished by first estimating patient-specific cell population similarity by performing pairwise Pearson correlations of the mean log-normalized expression profile of each cluster, then, high-similarity groups that included at least two clusters with r>0.986 were combined to produce an aggregate cluster. Since the number of effective expression profiles defining cluster centroids under this formulation is many magnitudes greater than the number of cell types in the dataset, merging cell populations with highly similar profiles allows molecular signatures from scRNA-seq profiles to indicate cell type identities. This merging operation and the removal of clusters with fewer than 50 cells reduced the cell-type space. Merged clusters were used to deconvolve RNA-seq profiles of patient samples, PDXs, and cell lines.

#### Cell type inference

We used the Celldex package v.2.0.0 in SingleR^41^ to determine the cell type of each population. To enable higher-resolution cell classification, including the distinction between multiple myeloid cell differentiation stages, we combined reference databases, including the Human Primary Cell Atlas^42^, Novershtern’s Hematopoietic Data^43^, Blueprint and ENCODE^44^, the Database of Immune Cell Expression^45^, and Monaco’s Sorted Human Immune Cells Database^46^.

#### Trajectory Inference Using Monocle 3

Trajectory analysis was conducted using the Monocle 3 package^47^ in R to infer the temporal progression of cellular states from single-cell RNA sequencing data. We first constructed a cell data set (CDS) object and applied the *preprocess_cds()* function, performing normalization using size factor normalization (norm_method = “size_only”), which accounts for differences in sequencing depth across cells. Dimensionality reduction was then performed via Principal Component Analysis (PCA) using the top 100 principal components (num_dim = 100) to capture major sources of transcriptional variance. The *reduce_dimension()* function was used to project the data into a two-dimensional manifold using Uniform Manifold Approximation and Projection (UMAP), preserving both local and global transcriptomic relationships. Unsupervised clustering of cells was performed using cluster_cells(), which constructs a k-nearest neighbor (kNN) graph and applies Louvain or Leiden community detection algorithms to identify transcriptionally distinct cell clusters. In addition to fine-grained clustering, Monocle 3 computes partitions—superclusters of communities—via pruning of the kNN graph, facilitating coarse grouping of cells across potentially broad biological states. These cluster and partition assignments were stored within the CDS object and later used in trajectory inference. Next, the *learn_graph()* function was used to infer a principal graph over the UMAP-reduced space using Reversed Graph Embedding (RGE). This method learns a smooth, branched graph structure that approximates the manifold of cell-state transitions. We set use_partition = FALSE to learn a unified graph across all cells, allowing for global trajectory inference rather than isolated trajectories within partitions. The graph learning process includes loop closure, which connects distal tips in the trajectory graph when their Euclidean and geodesic distances satisfy specified thresholds (defaults: euclidean_distance_ratio = 1, geodesic_distance_ratio = 1/3). Insignificant branches were pruned based on a minimal branch length threshold (default: 10 units), and tip nodes could optionally undergo orthogonal projection to refine placement (default: orthogonal_proj_tip = FALSE). The RGE algorithm constructs the graph by identifying latent paths that minimize distortion while preserving the topological structure of the reduced space, resulting in a biologically meaningful representation of lineage progression.

#### PDX-profile resolution

When analyzing bulk RNA-seq and scRNA-seq profiles of PDX samples, we classified the xenograft-derived sequence read data using *Xenome*^48^, which effectively handles a mix of reads from both the host and the graft, for further analyses.

#### CyTOF

Five COG diagnosis–relapse pairs profiled by scRNA-seq had a sufficient number of cells to perform CyTOF. After sorting, cells allocated for CyTOF were divided into two fractions; one was stimulated for 15 min with conditioned medium from HS-5 bone marrow stromal cells to activate intracellular signaling^49^, and the other was left unstimulated. Cells were fixed in 1.6% paraformaldehyde and barcoded (Cell-ID™ 20-Plex Pd Barcoding Kit, Fluidigm) for multiplexing, which allows combining cells for CyTOF analysis to reduce batch effects. Samples were then stained with 38 surface marker antibodies, permeabilized, and stained again for eight intracellular antigens; see Table S12 for the antibodies used. Normalization beads were included with every run. Doublets and non-viable (caspase 7^+^) cells were excluded. CyTOF assays were performed by the Baylor Cytometry and Cell Sorting Core. The data were compiled, cleaned, gated, and analyzed in Cytobank. Dimensionality reduction with UMAP was performed for each diagnosis– relapse pair using all surface and intracellular markers, except for cleaved caspase 7, which was used for the exclusion of non-viable cells.

#### GSEA to identify Expanded–Enriched Gene Sets and drivers of pAML transformation

Expanded Genes were identified by differential expression analysis comparing each expanded, relapsed, and stable pAMLs to all diminished pAMLs in each patient separately in the BCM cohort using Wilcoxon Rank–Sum tests. Genes expressed in at least 25% of the cells with an adjusted p<0.05 were considered significant for this comparison. Expanded Genes were defined as those significantly upregulated in at least 21 of our 52 comparisons. This cutoff was selected because it split the bimodal distribution of tallies across all candidate genes. We then used *enrichr*^50^ or hypeR^51^ with adjusted p<0.05 to identify gene sets and pathways that were enriched among Expanded Genes. We reverse-engineered a pAML-specific regulatory network based on TARGET pAML gene expression profiles as previously described^52–54^. This network was used to identify pAML-specific transcription factor (TF) targets that were used as gene sets for GSEA analysis. This, in turn, was used to estimate TF activity when comparing expression profiles across pAML aggregates.

#### Evaluating the predictive ability of models and gene sets

We used pAML clinical characterizations and RNA-seq profiles collected by TARGET and St. Jude Research Hospital—including clinical trials AAML1031, AAML0531, and AML08 to evaluate the prognostic predictive ability of gene sets and chemoresistant-cell models. The only treatment variability within this patient population was the combination of standard chemotherapy and bortezomib, but the results suggested that bortezomib did not affect overall survival.^7^ On the other hand, patients with high allelic ratio FLT3-ITD mutations were treated with sorafenib, which significantly affected patient outcomes. In total, TARGET profiles of AAML1031 included 1,330 diagnoses, 43 relapses, and 62 healthy donor bone marrow samples; TARGET profiles of AAML0531 patients included 144 samples with molecular and clinical annotations, whereas the St. Jude collection included 141 AML08 samples with molecular and clinical annotations. RNA-seq count data were log-normalized, and we performed GSVA^55^ analysis to evaluate whether gene sets were dysregulated in diagnosis and relapse samples (Figures S4 and S5). Comparisons of GSVA scores between samples from healthy donors, pAML diagnosis, and pAML relapse samples were evaluated by Wilcoxon Rank–Sum tests. GSVA scores were also used to evaluate the predictive ability of gene sets for patient outcomes. We next used SQUID^10^ to evaluate the abundance of pAML aggregates and non-pAML cell populations. Abundance estimates were evaluated for predictive ability for both OS and EFS by Cox regression with Bonferroni corrections for multiple testing. Outcome-predictive aggregates were composed of diagnosis sample cells and were predictive of both OS and EFS at an adjusted p<0.01. Tests of other cell populations yielded none that were outcome predictive. Because FLT3-ITD-positive patients were treated differently on the study’s protocol, we focused our analysis on FLT3–ITD– negative patients with profiled diagnosis samples.

In survival analyses, to limit machine learning, patients were equally partitioned based on the median inferred pAML aggregate abundance. Cumulative predictive aggregate abundance in the BCM and Lambo cohorts was combined using linear regression using AAML1031 patient profiles. The resulting parameters were used to analyze AAML0531 and AML08 data without additional learning. Survival analysis based on MRD after one or two rounds of chemotherapy split patients into MRD-positive vs. MRD-negative groups, and analysis based on cytogenetics assigned patients to one of three risk categories. Feature combinations partitioned the patient populations into discrete groups to predict total outcomes, including predictive aggregate abundance, MRD, and cytogenetics. In addition, we performed analyses using MRD after two rounds of chemotherapy and the AAML1831 risk-prediction algorithm. However, these variations did not alter the conclusions of our presented analysis based on cytogenetics biomarker aggregations and MRD alone; note that including MRD after two rounds of chemotherapy did not improve predictive accuracy, and that the AAML1831 risk-prediction algorithm is a refinement of the classic cytogenetics classification. Multivariate Cox proportional hazard models assessing pAML aggregate abundance, cytogenetics, MRD, and these factors combined were constructed using the *coxph* function in R.

When evaluating the dysregulation of transcription factor activity or regulatory networks immediately downstream from a transcription factor, we relied on the transcription factor’s inferred set of targets. Namely, we used the cumulative expression dysregulation of the targets of a transcription factor as a surrogate for the dysregulation of transcription factor activity (Figure S9 and supplementary Table S11).^23^ We also used the cumulative differential DNA accessibility of pAML signatures and transcription factor target sets based on Lambo et al. ATAC-seq profiles to evaluate regulatory modules that are epigenetically altered in pAML aggregates that acquire resistance.

#### PDX and cell line experiments and analysis

The sources of curated data from publicly available assays, including the targeting of PDXs and cell lines, as given in Figures 5 and S10, are provided in Table S13. When evaluating predictive aggregates, we required that every aggregate evaluated be present in at least 1% estimated abundance in control cells or animals. When evaluating the response of aggregates not predicted to be targeted, random samples of aggregates with sufficient estimated abundance were chosen. The size of random samples was set to match the number of predicted target aggregates, and sampling was repeated 10 times, with the geometric mean p-value reported in Figure S10. The median p-value random sample was selected for presentation in Figures 5 and S10.

Established serially transplanting pediatric AML PDX models from the PALeX resource were used for the experiments reported in this study (https://pdxportal.research.bcm.edu/pdxportal)^22^. Characteristics of patients from whom the models were derived and their viability after treatment with cytarabine and Saline are listed in Supplemental Table S8. For each experiment, 6–10 immunodeficient, human cytokine–producing mice (NSGS, MISTRG, or MISTRG6) were injected by tail vein with 2×10^5^ viable AML cells that were harvested from a prior passage and cryopreserved. Once mice demonstrated at least 2% human AML (hAML; human CD45^+^/CD3^‒^) cells in peripheral blood, they were assigned to receive cytarabine or saline (Figure S8). Treatment groups were balanced for peripheral blood disease burden. Cytarabine (50 mg/kg) was administered via intraperitoneal injection once daily for four days; control animals received an equal volume of saline.

All mice were humanely euthanized four days following the last dose of cytarabine or saline control (experiment Day 8). One mouse from each treatment group, for 6 of the 8 models that underwent cytarabine response evaluation, was profiled by scRNA-seq. Bone marrow and spleen were harvested, and the pAML proportion was determined by flow cytometry. Bone marrow from one cytarabine-treated and one saline-treated mouse was cryopreserved and profiled. We used diagonally weighted least squares estimation (DWLS)^56^ to estimate the number of cells matching outcome-predictive aggregates in each scRNA-seq cluster. We used GSVA to evaluate the enrichment of gene sets in PDX cluster pseudobulk profiles.

The effects of 1 µM treatment with all-trans retinoic acid (ATRA) in U937 and MOLM13 at 72 hours were estimated by Meier et al. with gene expression profiled by RNA-seq^57^. The effects of patient sample treatment by tamibarotene (100nM) at 24 hours were estimated by Perez et al., with gene expression profiled by RNA-seq^36^. We used SQUID to estimate the abundance of predictive aggregates in these samples, deconvolving RNA-seq profiles using all pAML and non-cancer aggregates as input. We note that the PDXs AML001 and AML006 were derived from patient samples p401 and p198, respectively. The correlation between predictive aggregate abundance and tumor variability was 0.95.

To test the effects of FLT3 and CDK4/6 inhibitors on chemoresistant pAML aggregates that were predicted to respond to FLT3 and CDK4/6 inhibition, we cultured cells from PDXs AML001, AML006, and AML903 in HS-5 conditioned medium and treated ex vivo with palbociclib, PF-07220060, or quizartinib (all 1 μM) or a combination of quizartinib and a CDK4/6 inhibitor for 3 days. Cells were harvested for flow cytometry determination of viability using Annexin V staining and differentiation using CD14 staining. Cells were also analyzed by bulk RNA-seq. We used DWLS to estimate the abundance of predictive aggregates in scRNA-seq profiles of saline-treated samples from these PDXs. The results suggested low, intermediate, and high cumulative abundance of predictive aggregates in AML006 (6%), AML903 (14%), and AML001 (24%). These estimates were significantly predictive of pAML tumor viability and CD14 expression in these PDXs. In particular, the correlations between predictive aggregate abundance and (1) tumor viability and (2) CD14 expression—a biomarker of myeloid differentiation—were 0.87 and 0.99, respectively.

To validate the effects of palbociclib and quizartinib in vivo, we treated NSGS mice (n=6) engrafted with AML903 with a combination of quizartinib (10 mg/kg) and palbociclib (100 mg/kg) or vehicle only (n=5) by daily oral gavage for 14 days, and followed the mice for the length of survival. Bone marrow was collected for bulk RNA-seq. When analyzing mouse cell lines, we used *biomaRt* package^58^ in R to retrieve human orthologs of mouse genes for making predictions about gene sets and to predict pAML aggregate abundance. Median survival of mice treated with palbociclib and quizartinib was longer (61 days) than that of mice treated with vehicle (29 days).

#### Evaluating potential patient response to inhibitors

When evaluating patients who may benefit from treatment with a given inhibitor (Figure S11), we first identified all predictive aggregates that exhibited a significant response to the inhibitor. Patients with high inferred abundance of these aggregates were then flagged as potential candidates for therapy with this inhibitor. To determine whether an aggregate’s response was significant, we generated a null distribution of responses using aggregates not predicted to be targeted. Predictive aggregates were considered significant if their response exceeded the null average by at least two standard deviations.

## AVAILABILITY OF DATA

TCH pAML scRNA-seq and bulk RNA-seq datasets, as well as the COG pAML scRNA-seq dataset, are available at GEO Super Series GSE271137 (reviewer token kjolckuqllehbwl). Bulk RNA-seq data for the FLT3 inhibitor quizartinib and CDK4/6 inhibitor palbociclib treatments are also included in the same GEO accession. The COG pAML CyTOF profile data reported in this paper are publicly available via Cyto-bank. Note that the expression profiles of 65 protein markers were included for 12 of the TCH pAML samples, but these were not used in this study.

## ACKNOWLEDGEMENTS

We thank Sridevi Addanki, Maci Terrell, Raushan Rashid, Jaden Sherman, Alan Gonzalez, Hailey Oviedo, Noah Keogh, Julia Kim, Ritesh Dontula, Tamilini Ilangovan, Rehan Siddiqui, and Michelle Alozie for help with the implementation of animal experiments. This project was partially funded by Wipe Out Kids Cancer, the Helis Medical Research Foundation’s Helis Research Program Grant, CPRIT awards RP240432, RP180672, RP180674, RP200504, RP170691, RP220646, and RP230120; European Union’s Horizon 2020 research and innovation program under grant agreement 826121; and the NIH awards U24CA196173, U24CA114766, U10CA180899, U10CA180886, U24CA196173, R21CA223140, R21CA286257, CA125123, RR024574, OD036336, and OD038251. Baylor College of Medicine Advanced Cores are supported by NIH grants P01CA261669, S10OD018033, S10OD023469, S10OD025240, and P30EY002520. This project was supported by the Cytometry and Cell Sorting Core at Baylor College of Medicine, and the assistance of Joel M. Sederstrom. The COG Biospecimen Bank and PDX development were also supported by funding from the St. Baldrick’s Foundation U24CA196173, Texas Children’s Hospital Pediatric Pilot Research Fund, CURE Childhood Cancer, Leukemia and Lymphoma Society/Blood Cancer United, Target Pediatric AML/the Children’s Oncology Group Foundation, and gifts of funding from the Turn it Gold Fund. Support from the Cytometry and Cell Sorting Core includes NIH P30 AI036211, P30 CA125123, and S10 RR024574. The results published here are wholly or in part based upon data generated by the TARGET initiative, phs000218.v8.p1, managed by the NCI. The data used for this study is available through Genomic Data Commons. The content is solely the responsibility of the authors and does not necessarily represent the official views of the National Institutes of Health. This manuscript was prepared with the assistance of a science writer, Ariel M Lyons-Warren, MD, PhD.

## AUTHORSHIP

M.J.N., T.K.M., M.S.R., and P.S. designed the research and wrote the paper. A.M. Lyons-Warren managed, edited, and contributed to the writing of the manuscript. A.M.S., M.J.K., M.R., S.S., L.K., and J.S.Y. performed experiments. H-S.C., J.E., B.Z., M.R.M., M.R., M.J.N., S.U., B.B., M.S.R., and P.S. analyzed data.

Conflict-of-interest disclosure: The authors declare no competing financial interests.

## SUPPLEMENTARY FIGURES

**Figure S1.**
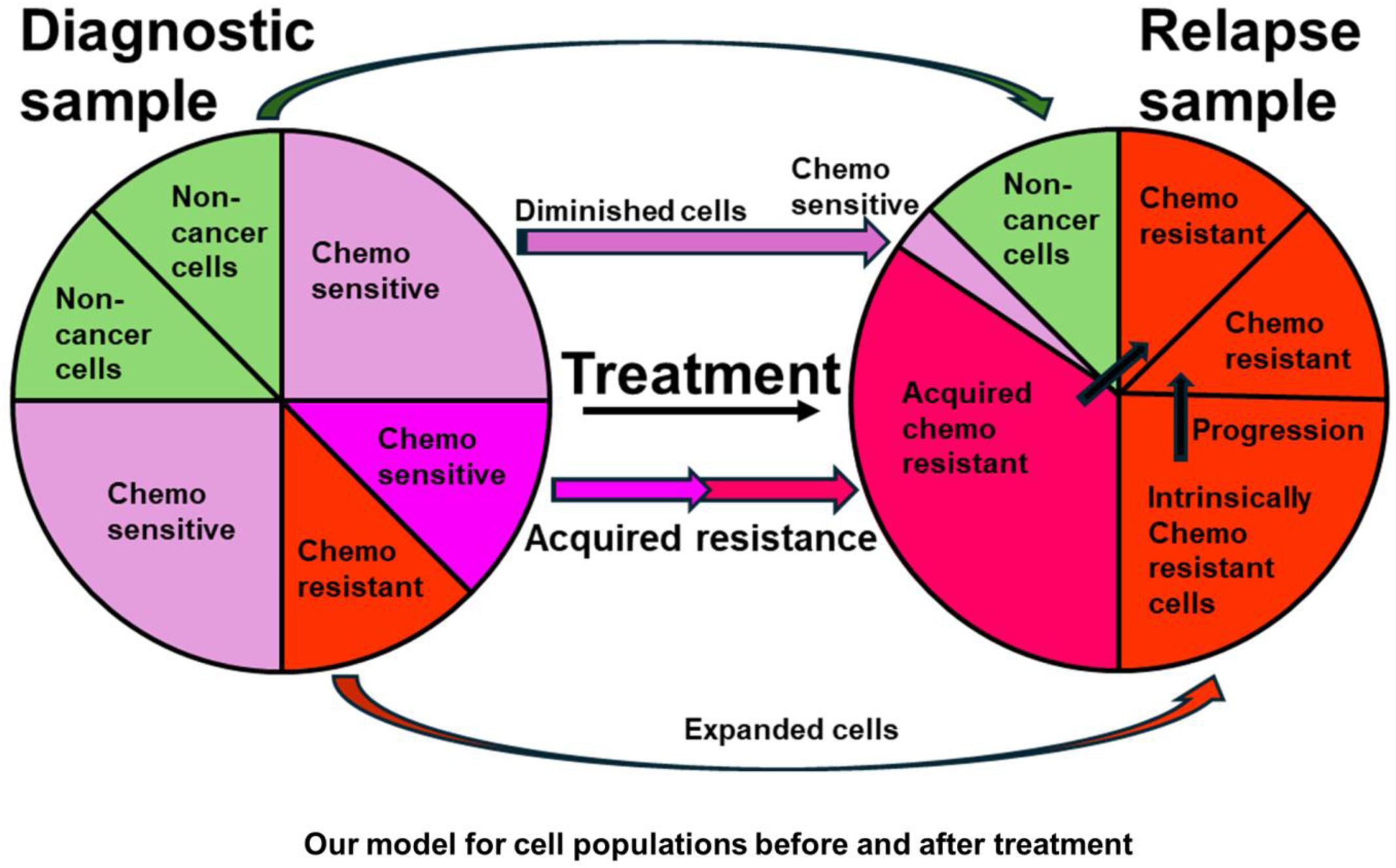
Our proposed model for the cell composition of pAML patient samples at diagnosis and relapse accounts for non-cancer cell types and both chemoresistant and chemosensitive pAML cells in the diagnosis sample. This model proposes the following: **1**) the relative proportion of non-cancer cell types in the sample may increase or decrease between diagnosis and relapse, **2**) the relative proportion of chemoresistant cells at diagnosis (expanded pAMLs) will increase during treatment, and **3**) the relative proportion of some chemosensitive cells will diminish (diminished pAMLs), whereas others that display transformed transcriptomics may gain resistance to chemotherapy. We aim to characterize expanded and transforming cells that are predictive of pAML patient outcomes.

**Figure S2.**
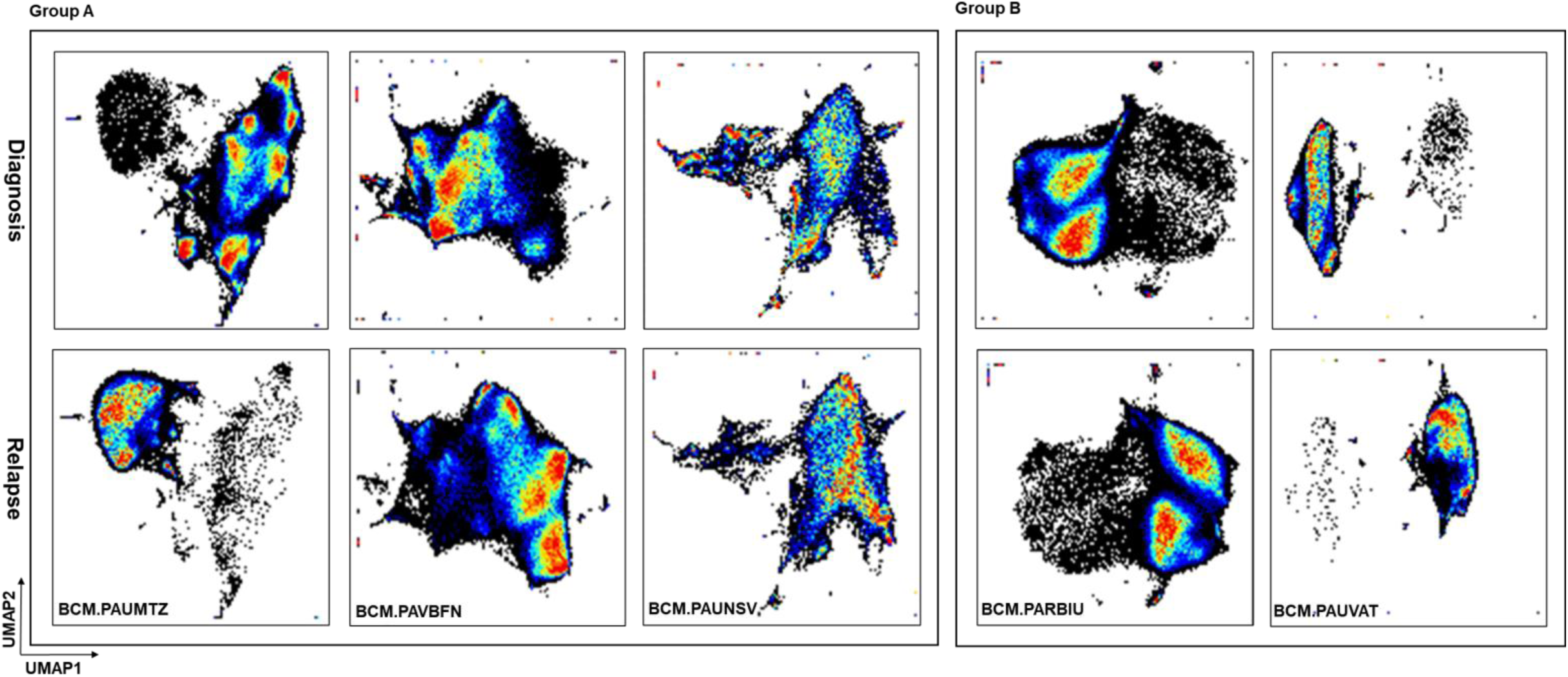
UMAP visualization of cytometry by time of flight (CyTOF) profiles of the five representative pAML pairs from Group A and B patients, shown in Figure 2D. UMAP color gradients represent cell density.

**Figure S3.**
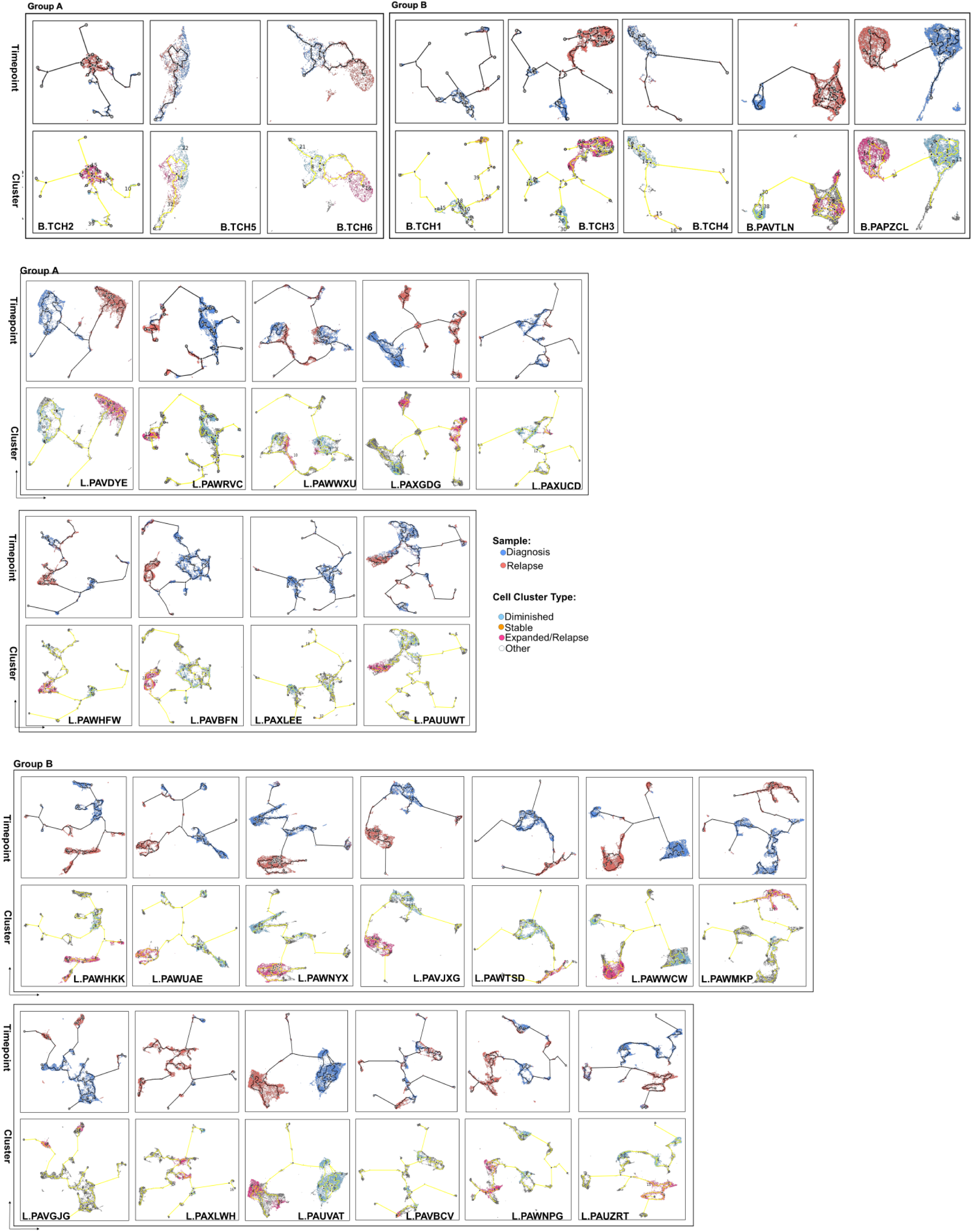
Single-cell trajectory plots, analogous to Figure 2D, depict profiles of samples from patients with or without expanded pAMLs in diagnostic samples (i.e., Group A and Group B for BCM and Lambo cohort patients). In the top panel, diagnosis and relapse time points are color-coded in red and blue, respectively, with the inferred trajectory path indicated by a solid black line. In the bottom panel, clusters are categorized as Expanded (in light red), Stable (orange), Diminished (in light blue), or Other (in white), with the inferred trajectory path represented by a solid yellow line. Trajectory paths predict cell transformations.

**Figure S4.**
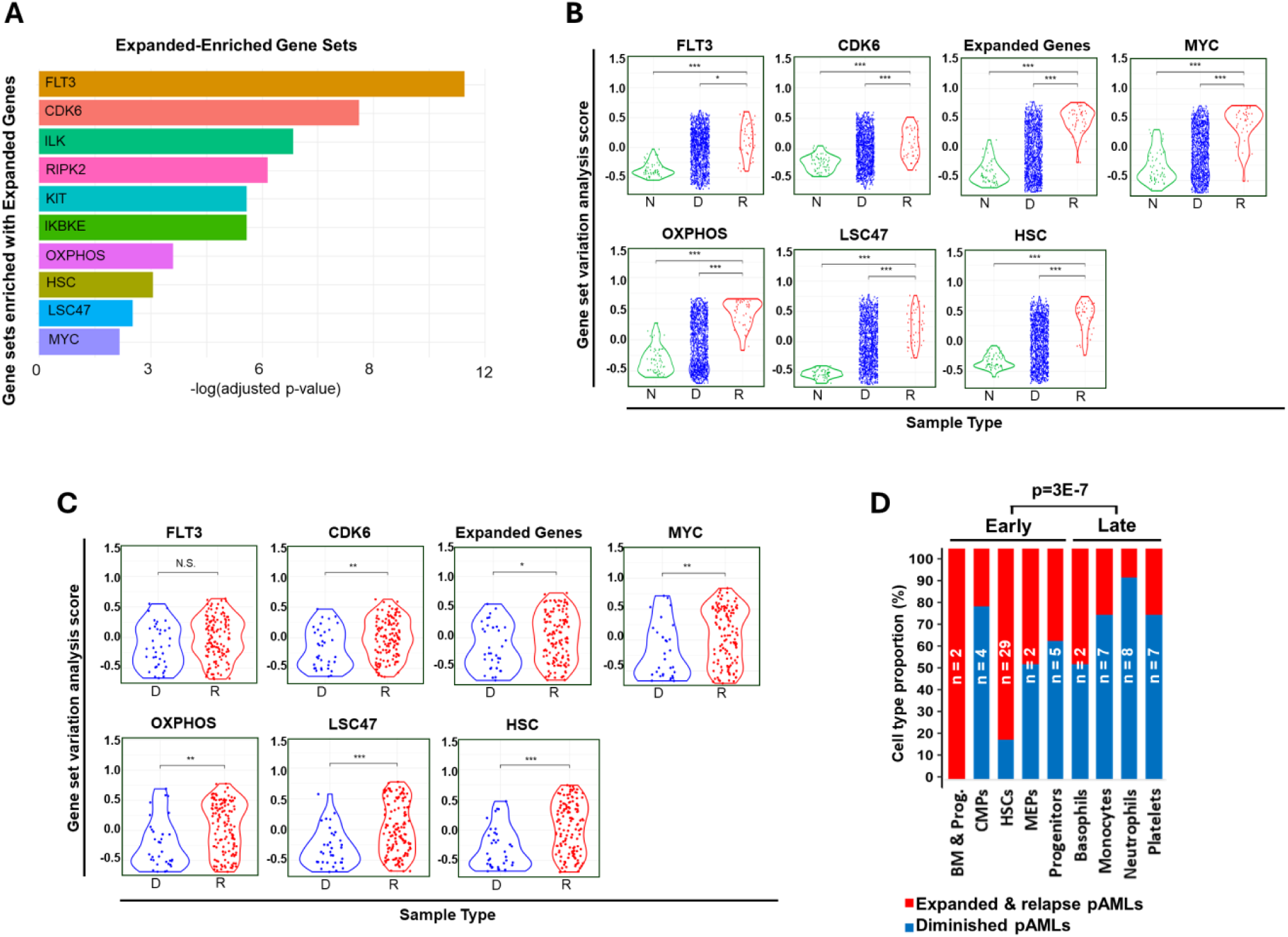
(A) Gene sets and pathways that were significantly enriched with Expanded Genes (Expanded–Enriched Gene Sets), including genes that are co-expressed with Fms-related receptor tyrosine kinase 3 (FLT3), cyclin-dependent kinase 6 (CDK6), integrin-linked kinase (ILK), receptor-interacting serine/threonine kinase 1 (RIPK), KIT proto-oncogene (KIT), and inhibitor of nuclear factor kappa-B subunit epison (IKBKE), as well as pathways including Hallmark oxidative phosphorylation (OXPHOS), hematopoietic stem cell (HSC), leukemia stem cell populations gene set (LSC47), and MYC targets. **(B, C)** Gene Set Variation Analysis (GSVA) revealed Expanded– Enriched Gene Sets that are upregulated in (B) AAML1031 and (C) AML08 relapse samples; N, bone marrow aspirates from non-cancer donors; D, diagnosis samples; R, relapse samples. **(D)** The proportion of expanded and diminished pAML cell clusters in the BCM cohort as a function of pAML differentiation showed an enrichment of expanded and relapse pAML cell clusters in undifferentiated pAML aggregates.

**Figure S5.**
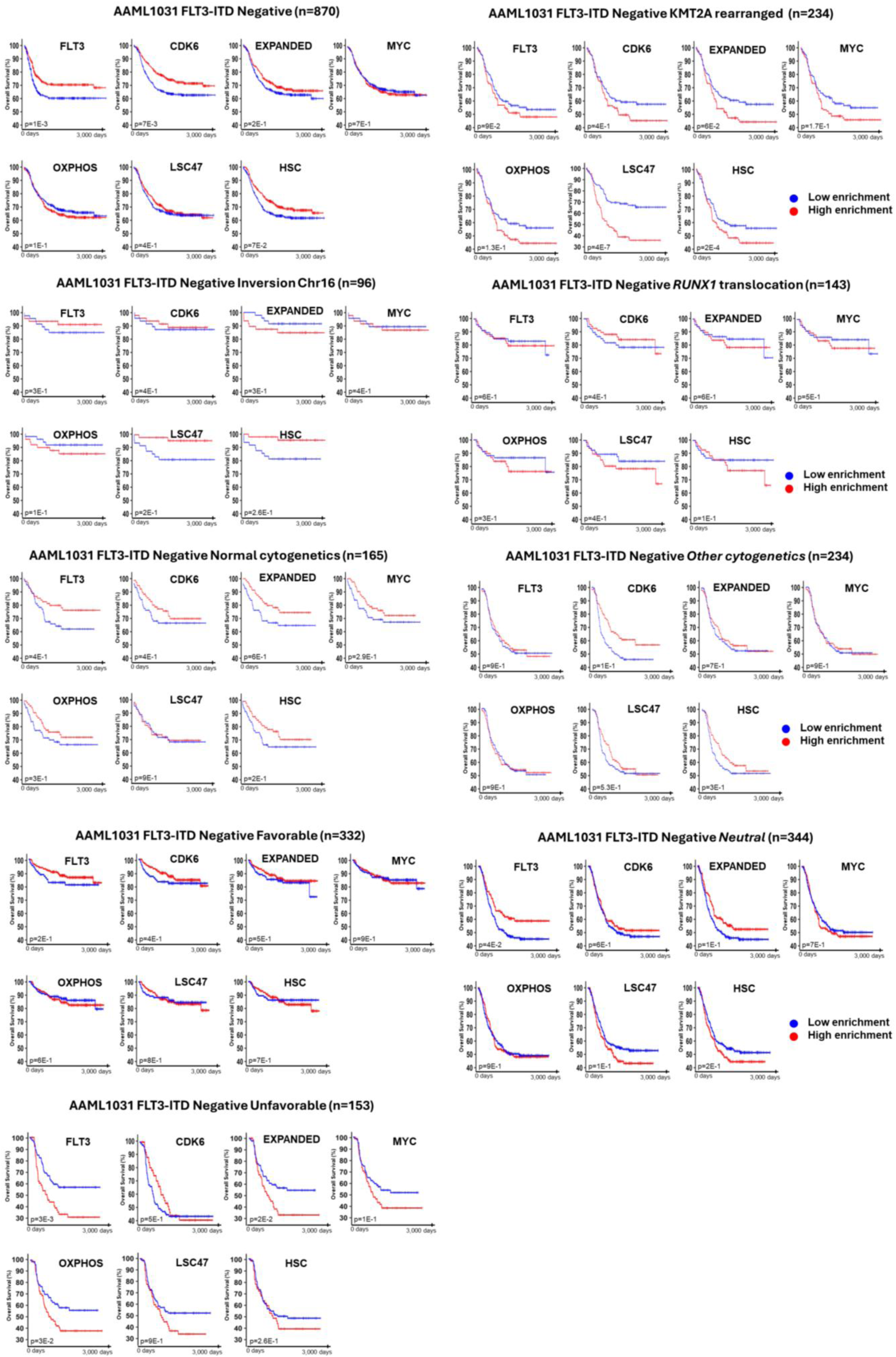
GSVA of Expanded–Enriched Gene Sets in cytogenetic classifications of AAML1031 without correction for multiple testing. We evaluated the outcomes-predictive power of previously proposed outcomes-predictive pathways and gene sets, including LSC47, HSC, and OXPHOS, as well as genes that were recurrently dysregulated in chemoresistant pAML aggregates (*Expanded*). We also evaluated whether MSigDB-curated pathways were predictive of outcomes, and identified genes co-expressed with FLT3 and CDK6 and the MYC pathway as the most predictive. After multiple testing corrections, the following were identified as significantly predictive: LSC47 and HSC enrichment were significant in KMT2A rearranged pAML, but not in the Unfavorable Cytogenetics category or any other categories. Expanded Genes were enriched in the Unfavorable Cytogenetics category (p=2E-2).

**Figure S6.**
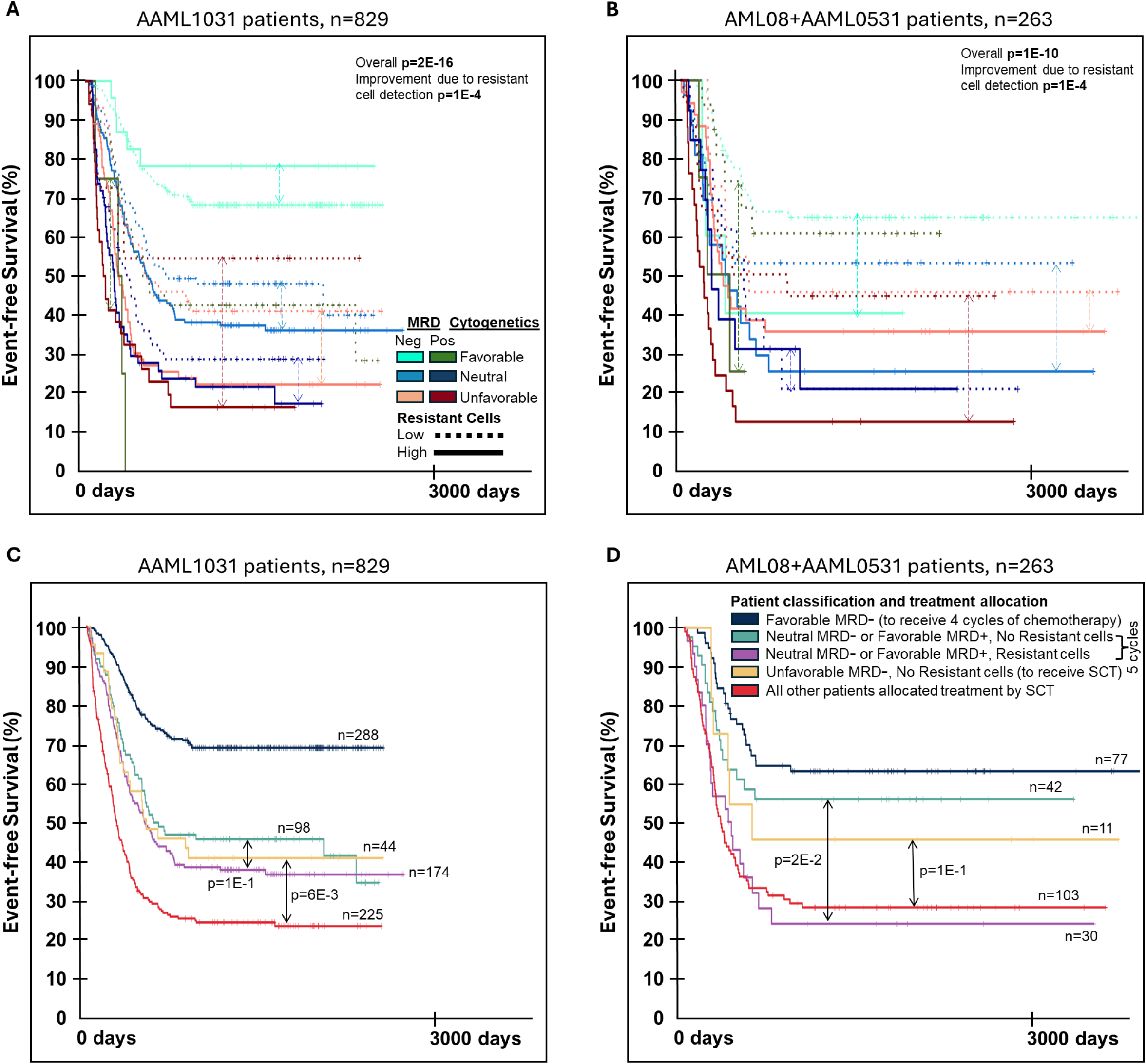
EFS analysis of predictive aggregates. Analogous to Figure 3. (**A–B**) Estimated abundance of chemoresistant pAML aggregates (resistant cells) significantly predicted EFS for patients in (A) AAML1031 and (B) AML08 + AAML0531 cohorts, as determined by Kaplan–Meier survival analysis. This included patients classified by MRD after one chemotherapy cycle and stratified by cytogenetics (Favorable, Neutral, Unfavorable) according to the COG AAML1831 protocol. Across independent trials, resistant cell abundance improved EFS prediction (p = 1E-4). Solid and dotted lines represent high-and low-abundance patients, respectively. Arrows indicate patients within the same AAML1831 risk categories. (**C–D**) Resistant cell abundance reclassified patients in (C) AAML1031 and (D) AML08 + AAML0531 who were allocated identical therapies under the current AAML1831 protocol. This analysis revealed higher-risk patients (purple) who would not be prescribed SCT under the COG AAML1831 protocol, with EFS < 0.4, and lower-risk patients (orange) allocated SCT under AAML1831 but achieving EFS > 0.4 without SCT across AAML1031 and AML08 + AAML0531.

**Figure S7.**
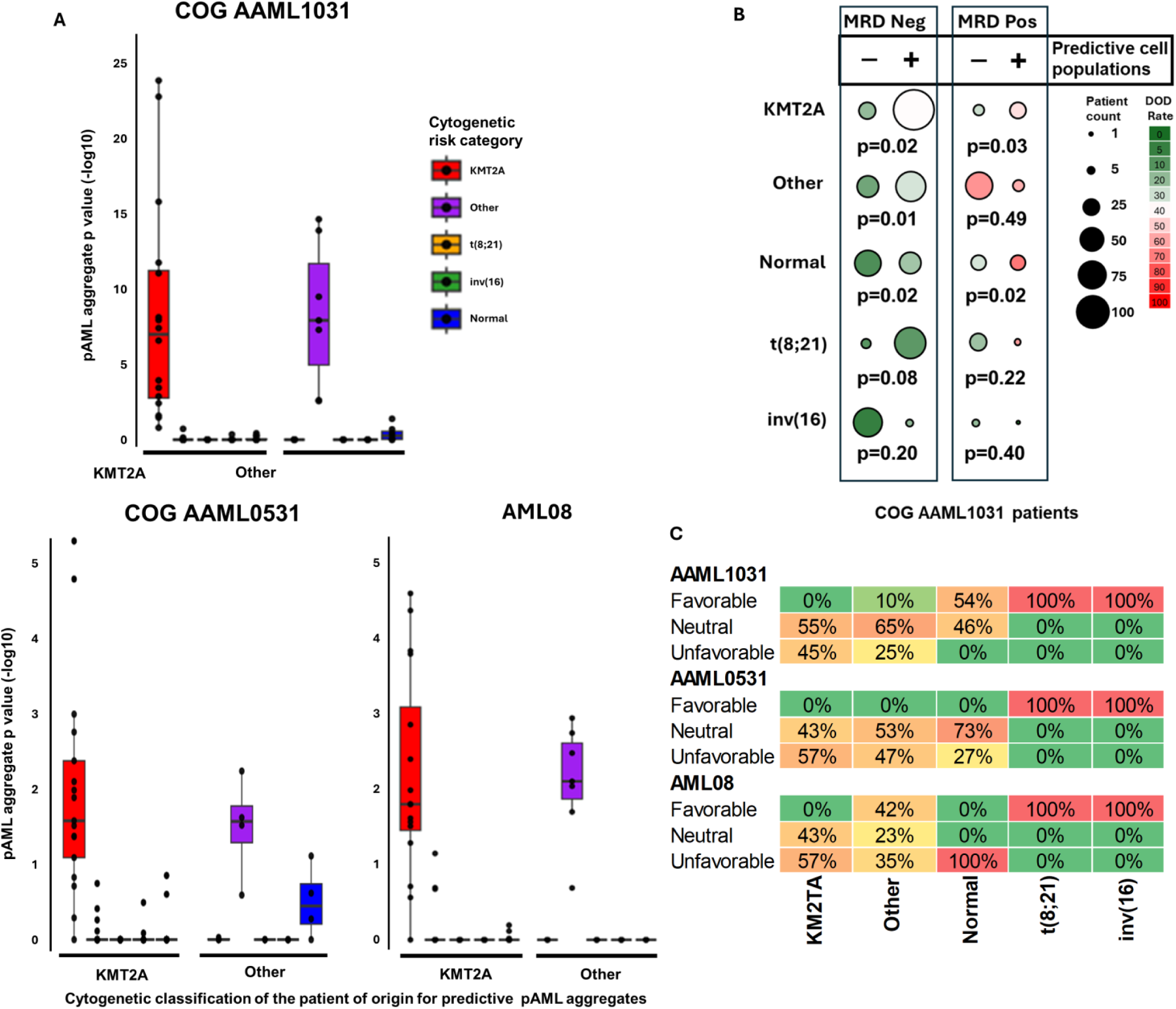
(A) We classified predictive pAML aggregates by the classic cytogenetic classifications of the patients that they were derived from, e.g., KMT2A predictive aggregates were derived from a pAML tumor with KMT2A rearrangements. Then, we classified AAML1031, AAML0531, and AML08 patients into the same cytogenetic classifications, producing patient classifications. Finally, for each pAML aggregate classification, we evaluated whether the estimated abundance of pAML aggregates from this classification is significantly greater in each patient classification relative to others. Our analysis revealed that the abundances of predictive aggregates from one classification is significantly greater in patients with the same classification, i.e., predictive aggregates with KMT2A alterations were significantly more abundant in patients with KMT2A alterations, etc. However, predictive aggregates were also identified and were predictive in patients who did not match their classification. (**B**) Focusing on AAML1031 patients, predictive aggregates were significantly predictive of outcomes in KMT2A, Other, and Normal cytogenetic classifications. Marker size denotes the number of patients from each cytogenetic classification with detected predictive aggregates, and the color denotes the rate of poor outcomes. (**C**) A mapping between the cytogenetic classification used here {KMT2A, Other, Normal, inv(16), and t(8;21)} and the risk categories proposed in the COG AAML1831 trial {Favorable, Neutral, and Unfavorable}. The detection of inv(16) or t(8;21) was consistently associated with low-risk, favorable prognosis; however, risk in other categories has been refined by COG AAML1831. DOD: died of disease.

**Figure S8.**
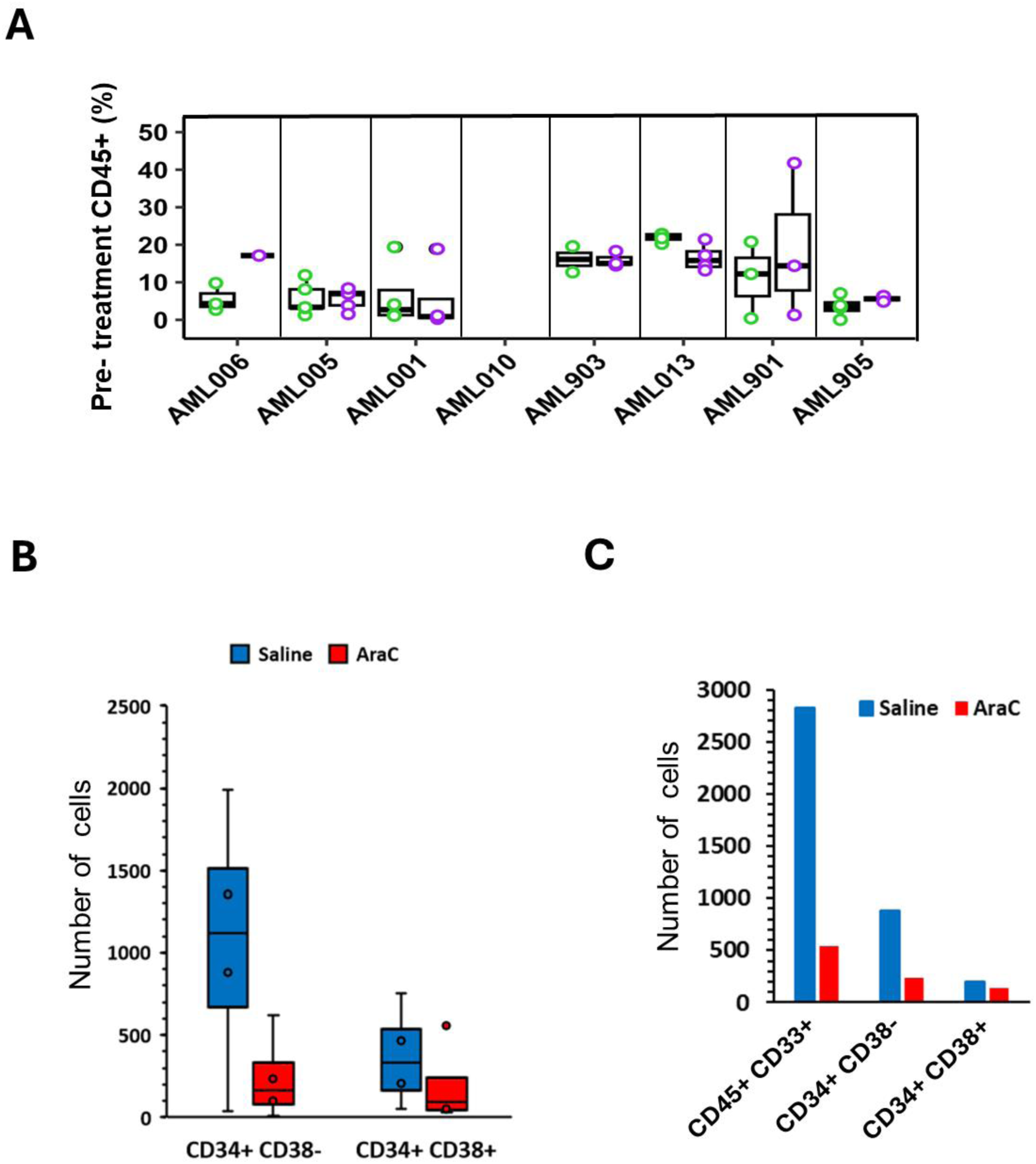
(A) Pre-treatment CD45^+^ cell abundance in PDX samples was used to determine the treatment initiation time point. (**B**) Graphs showing the number of CD34^+^CD38^‒^ and CD34^+^CD38^+^ pAML cells among total residual human CD45^+^CD33^+^ cells identified in patient-derived xenograft (PDX) models AML006, AML005, AML001, and AML010 after treatment with cytarabine (AraC; red bars) or saline (blue bars). (**C**) Total CD45^+^CD33^+^, CD34^+^CD38^‒^, and CD34^+^CD38^+^ pAML cells in PDX AML006 after treatment with AraC (red bars) or saline (blue bars).

**Figure S9.**
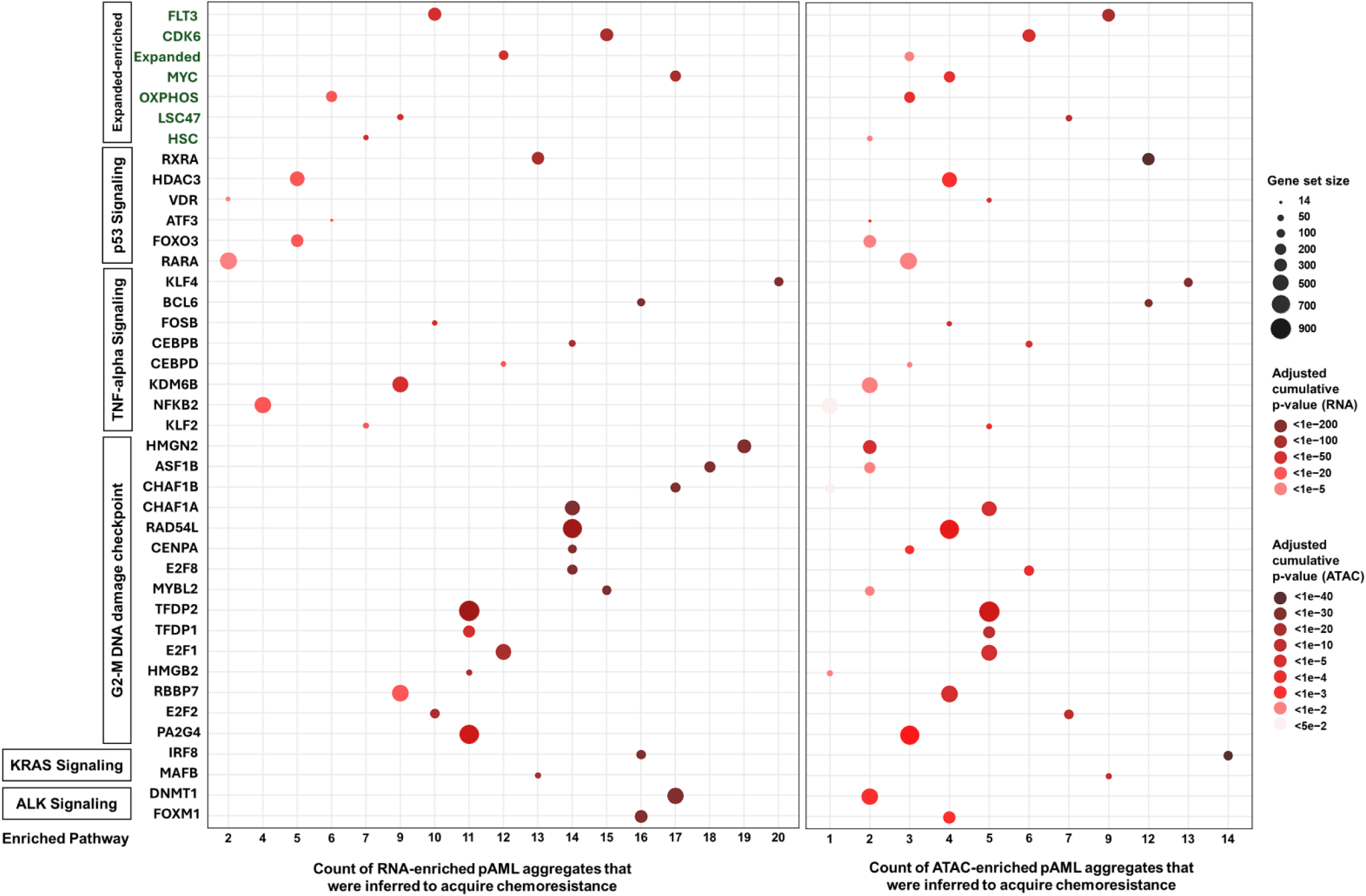
Gene sets and regulatory network modules that were enriched with differential gene expression (left) and chromatin accessibility (right) in pAML aggregates that were inferred to acquire chemoresistance, relative to pAML aggregates that were inferred to be chemosensitive in the same sample. The total number of pAML aggregates that were inferred to acquire chemoresistance is shown on the x-axis. Gene sets and regulatory network modules anchored at specific transcription factors are shown on the y-axis. The dot size and color denote the number of genes and the p-value for the enrichment (adjusted and cumulative over pAML aggregates).

**Figure S10.**
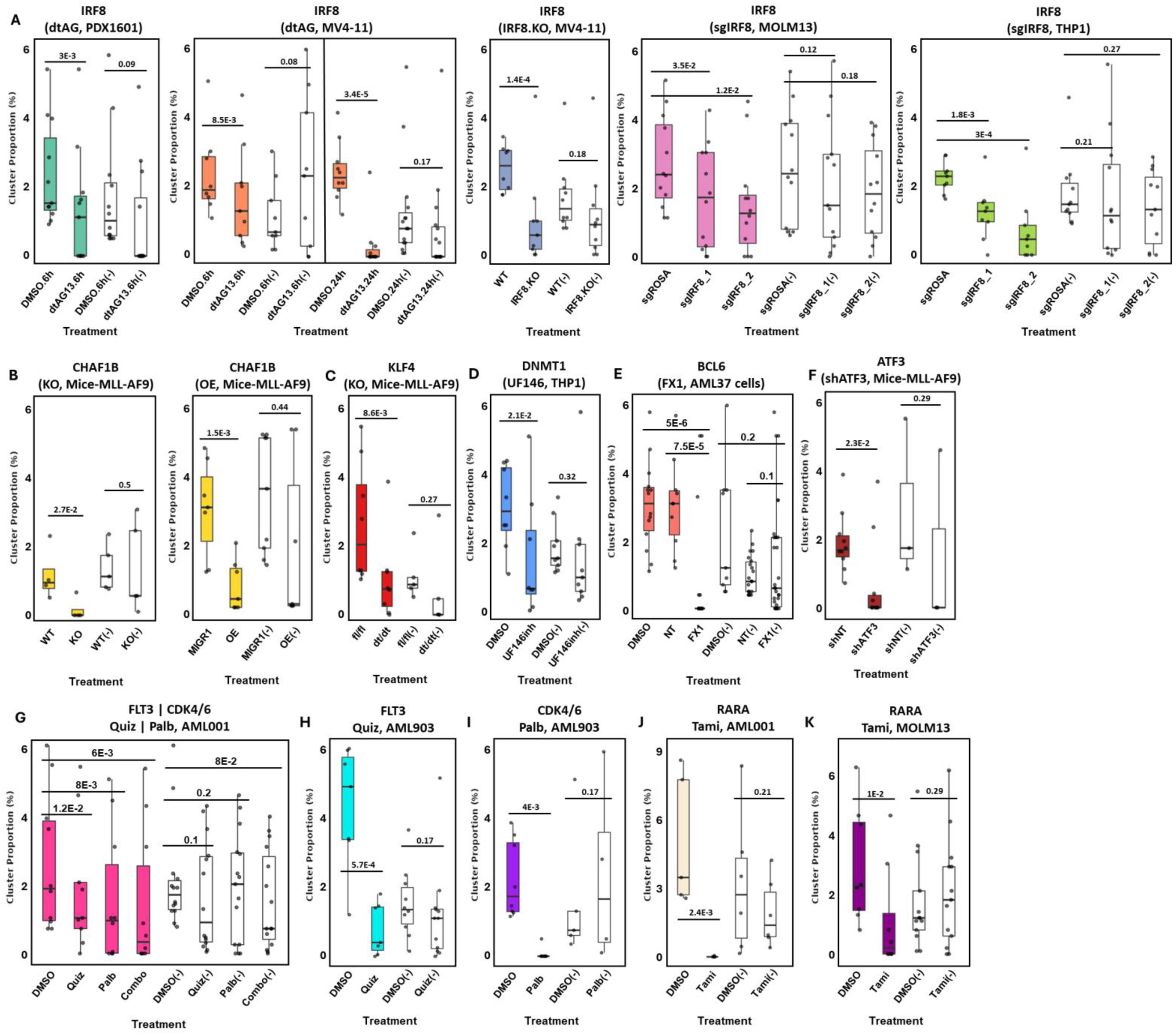
Specific evaluation of gene sets and regulatory network modules that were enriched with dysregulated genes and differential chromatin accessibility in diminished pAMLs that were inferred to acquire chemoresistance. We compared the inferred abundance of pAML aggregates (1) with predicted vulnerabilities in cells and tumors, and (2) those that were not predicted to be vulnerable to the effects of specific targeting. These results include a detailed comparison to the response of pAML aggregates not predicted to be targets corresponding to Figure 5L. The abundance of pAML aggregates that were not predicted to be vulnerable to each perturbation is shown in white for each gene set and regulatory network module. The figures present random selections of pAML aggregates to match the number of detected pAML aggregates that were predicted to respond to each perturbation.

**Figure S11.**
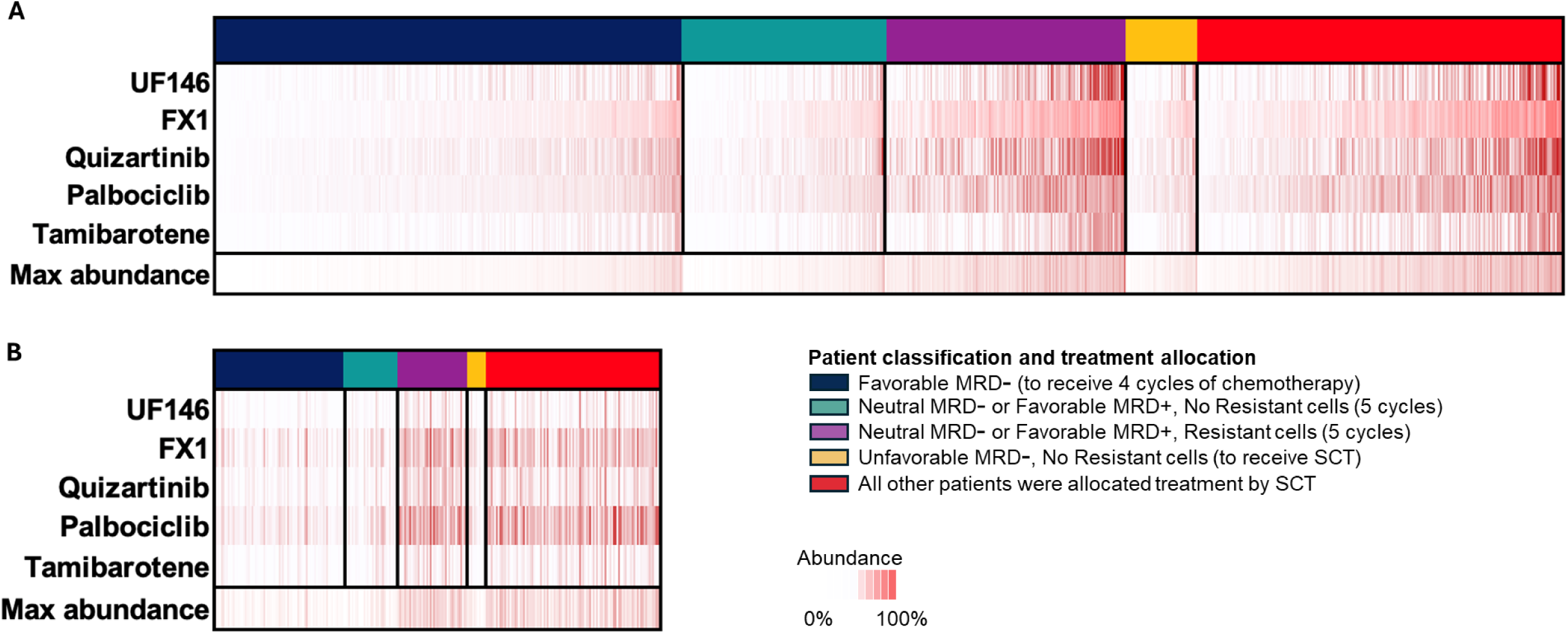
(**A**) AAML1031 and (**B**) AAML0531+AML08 patient samples predicted to contain pAML aggregates that respond to the five tested inhibitors in Figures 5 and S10. Namely, for each inhibitor, we collected the pAML aggregates whose abundance was significantly reduced, with significance estimated based on a null composed of non-target pAML aggregates (at least two standard deviations away). For each diagnostic patient sample, we evaluated the total inferred composition of these pAML aggregates. Color denotes the total proportion (from 0 to 1).

## SUPPLEMENTARY TABLES

**Table S1.**Clinical annotation of each profiled pediatric acute myeloid leukemia (pAML) patient in the BCM and Lambo cohorts, including age, sex, cytogenetics, mutations, time to relapse, measurable residual disease (MRD), and sample blast abundance.

**Table S2.**The table form of Figure 1C shows cluster frequencies in each profiled sample.

**Table S3.**Cluster frequency tables corresponding to Figures 2A,B for each analyzed patient. Cell counts by sample are provided. Each tab provides cell counts per cluster where the profiles of a patient sample and 3 normal bone marrow samples were merged.

**Table S4.**Total cell counts and predicted cell types for each cluster are reported in Table S3.

**Table S5.**Differentially expressed genes and the statistics used for each comparison in Figures 2A,B.

**Table S6.**The identity of our Expanded–Enriched Gene Signatures.

**Table S7.**MRD, cytogenetic classification, abundance of predictive aggregates, EFS, and OS status of patients in the three clinical trials evaluated.

**Table S8.**A description of PDX models and response to therapies corresponding to Figures 4A,B.

**Table S9.**Cell counts in clusters identified in scRNA-seq PDX profiles (Figure 4D).

**Table S10.**Inferred abundance of pAML aggregates in each PDX cluster.

**Table S11.**Inferred transcription factor activity and gene set enrichment (by GSEA) in pAML aggregates that were inferred to acquire chemoresistance. Both gene expression and DNA accessibility enrichment are provided. Gene sets used are described; p-values were Bonferroni corrected.

**Table S12.**The antibodies that were used to analyze the five pAML profiles using cytometry by time of flight (CyTOF). Both raw and normalized data were deposited in Cytobank.

**Table S13.**Reference to the assays and studies used to evaluate pAML aggregate vulnerabilities in Figures 5 and S10.

## Notes

### Competing Interest Statement

The authors have declared no competing interest.

### Summary of Updates

New results included and figures fixed.

